# Identification of key interactions of benzimidazole resistance-associated amino acid mutations in *Ascaris* β-tubulins by molecular docking simulations

**DOI:** 10.1101/2021.11.03.467184

**Authors:** Ben P. Jones, Arnoud H.M. van Vliet, E. James LaCourse, Martha Betson

## Abstract

*Ascaris* species are soil-transmitted helminths that infect humans and livestock mainly in low and middle-income countries. Benzimidazole (BZ) class drugs have predominated for many years in the treatment of *Ascaris* infections, but persistent use of BZs has already led to widespread resistance in other nematodes, and treatment failure is emerging for *Ascaris*. Benzimidazoles act by binding to β-tubulin proteins and destabilising microtubules. Three mutations in the β-tubulin protein family are associated with BZ resistance. Seven shared β-tubulin isotypes were identified in *Ascaris lumbricoides* and *A. suum* genomes. Benzimidazoles were predicted to bind to all β-tubulin isotypes using *in silico* docking, demonstrating that the selectivity of BZs to interact with one or two β-tubulin isotypes is likely the result of isotype expression levels affecting the frequency of interaction. *Ascaris* β-tubulin isotype A clusters with helminth β-tubulins previously shown to interact with BZ. Molecular dynamics simulations using β-tubulin isotype A highlighted the key role of amino acid E198 in BZ-β-tubulin interactions. Simulations indicated that mutations at amino acids E198A and F200Y alter binding of BZ, whereas there was no obvious effect of the F167Y mutation. In conclusion, the key interactions vital for BZ binding with β-tubulins have been identified and show how mutations can lead to resistance in nematodes.

## Introduction

The large intestinal roundworm *Ascaris lumbricoides* infects humans and causes ascariasis. *Ascaris lumbricoides* is a parasitic nematode that resides in the small intestine of its host and can persist there for up to 2 years ^1^. Ascariasis is often asymptomatic, but in regions of high *A. lumbricoides* prevalence there can be significant effects on host wellbeing, with chronic ascariasis leading to reduced cognitive ability and stunted growth due to malnutrition ^2^. The migrating larvae may also cause pulmonary ascariasis which results in asthma-like symptoms, whilst high worm burdens can lead to more serious pathologies such as organ blockages, which can result in death ^3,4^. As of 2019 there was an estimated 446,000 people infected with *A. lumbricoides* worldwide with an estimated loss of 754,000 disability adjusted life years (DALYs) ^5^. Most of these infections occur in rural and poor urban areas of low- and middle-income countries, where hygiene and sanitation infrastructure can be of a lower standard than in higher income areas, and therefore people are more exposed to infection. *Ascaris suum* is a closely related roundworm of pigs, although it can also be zoonotic ^6,7^. *Ascaris suum* has a wider geographical distribution than *A. lumbricoides* and is one of the most prevalent intestinal parasites of pigs worldwide ^8,9^. *Ascaris suum* infection can lead to production losses from reduced growth rates, altered muscle composition and the condemnation of livers due to fibrotic lesions known as milk spots ^10,11^.

There are only a small number of drugs available to treat ascariasis, which include the benzimidazoles (BZ), macrocyclic lactones and levamisole ^12^. Overreliance on these drugs has led to the potential for drug resistance. Mass drug administration (MDA) of BZ anthelmintics, such as albendazole and mebendazole, in endemic regions is the strategy for control and elimination of a number of helminth diseases in humans, including ascariasis. The most recent 2021-World Health Organization roadmap for neglected tropical diseases has targeted the elimination of ascariasis as a public health problem in 96 countries by reaching 75% coverage of MDA in targeted populations ^13^. Whilst repeated treatment in endemic communities may be able to reduce parasite burdens, it does not prevent reinfection, and it is well-established that pressure applied by MDA can lead to the evolution of drug resistance ^14^. Benzimidazole resistance has been detected in many intestinal parasites of both veterinary and human importance, and the first signs of reduced susceptibility in *Ascaris* have been reported ^15–24^. To date, BZ resistance has been linked to mutations in β-tubulin proteins, more specifically at amino acids 167, 198 or 200, based on the *Haemonchus contortus* β-tubulin reference sequence (accession number: AAA29170.1). Nematodes usually encode multiple β-tubulin isotypes but not all are expressed equally, with some being life-stage or cell-type specific ^25^. One of the highly expressed isotypes, β-tubulin isotype 1, is commonly linked to resistance in parasitic nematodes ^16,18–21,26–29^. Little is known about the contribution of other β-tubulin isotypes to drug interactions and resistance. Based on evidence *from Caenorhabditis elegans*, it is likely that most of these isotypes are redundant or have specialised roles within specific cells or at certain developmental stages ^30^. So far, no work has been done to characterise the roles of the β-tubulins in ascarids or other common STHs and therefore the role they play in drug mechanisms and the development of BZ resistance is still unknown.

One of the biggest hindrances to answering these questions is the ability to culture the full lifecycle of these parasites *in vitro*, as well as the ethical considerations and costs associated with studying parasites in animal models. *In silico* approaches could help to solve these problems by predicting the differences seen between proteins and how they interact with drugs. *In silico* docking is a technique that uses computational software to try and mimic biological systems and monitor molecular interactions. A common use is to model protein-ligand docking to theoretically assess the ability of a ligand to bind within the active sites of a protein and to develop novel drugs ^31^. *In silico* docking has been performed using the β-tubulins of several helminths including *H. contortus, Trichinella spiralis* and filarial nematodes ^32–35^. These studies have highlighted the changes in protein conformation that occur when resistance mutations are present and how that affects drug interactions. To date these methods have not been applied to *Ascaris*, nor has any study looked into the differences that may be seen between the individual β-tubulin isotypes within a genus or species.

The aims of this study were to investigate the interactions between commonly used BZ drugs and *Ascaris* β-tubulins, and identify what changes occur when mutations are present. The first objective was to confirm that BZ binding in *Ascaris*. was similar to that of other helminths. The second objective was to compare the binding of these drugs in each of the β-tubulin isotypes present in *Ascaris*. The final objective was to repeat these experiments in proteins that contain the common resistance-associated mutations, to gain an insight into changes that lead to resistance.

## Results

### Identification of *Ascaris* β-tubulin isotypes

Twenty-one β-tubulin sequences were retrieved from NCBI, which were reduced to six after removal of partial and duplicated sequences. BLAST searches against the three *Ascaris* genomes in Wormbase-Parasite identified a total of 122 matches, 51 of which were β-tubulins based on nomenclature and identity to reference sequence, with the remainder α-tubulins. Of the 51 β-tubulins, after duplicates had been removed, there were 8 sequences remaining from the *A. lumbricoides* genome (GCA_000951055.1), seven from one *A. suum* genome (GCA_000298755.1) and six from a second *A. suum* genome (GCA_000187025.3). To extend the search for more distantly related tubulins, the Exonerate program predicted the presence several additional tubulin sequences in the three *Ascaris* genomes. A search of the Conserved Domain Database revealed that most were α-tubulins, but a new β-tubulin sequence was identified for *A. suum* (E’) and a more complete sequence for *A. lumbricoides* B’ sequence was identified and added to the sequences used for phylogenetic analysis (Supplementary Table S1). The isotype G identified from *A. suum* (GCA_000187025.3) was found to be split into two consecutive genes in the genome annotation, although manual alignment of these two genes with isotype G from *A. lumbricoides* confirmed that these two genes represented two halves of the full gene with an incorrect stop codon predicted at the end of an exon (at position 169-171 of the cDNA). Therefore, these two consecutive genes were concatenated and used as the *A. suum* isotype G gene for all further work (Supplementary Fig S1). The protein to gene alignment undertaken with the Exonerate program on the two newly available genomes (*A. lumbricoides* GCA_015227635.1 and *A. suum* GCA_013433145.1) found sequences for all isotypes, with the exception of isotype G in *A. lumbricoides*. These sequences were added to the existing data and a phylogeny was created which included β-tubulins from other Ascaridomorpha species. A full list of sequences used can found in Supplementary Tables 1 and 2.

The phylogenetic tree showed a clear separation into definitive isotypes that appear to have diverged early in the evolution of the Ascaridomorpha infraorder (Fig. 1). When phylogenetic trees for the amino acid and nucleotide sequences were compared, the structuring of the isotype clades were not consistent, although similar relationships were observed between sequences within clades. *Ascaris suum* isotype F3 had one truncated exon and so did not fit into the group as well as other sequences. *Ascaris suum* isotype E’ was also seen to be divergent from the rest of the isotype E group, and as this sequence was only found in one genome it was not designated its own isotype. The effects of these sequence variations were seen more clearly in the phylogenetic tree based on amino acid sequences.

**Figure 1:**
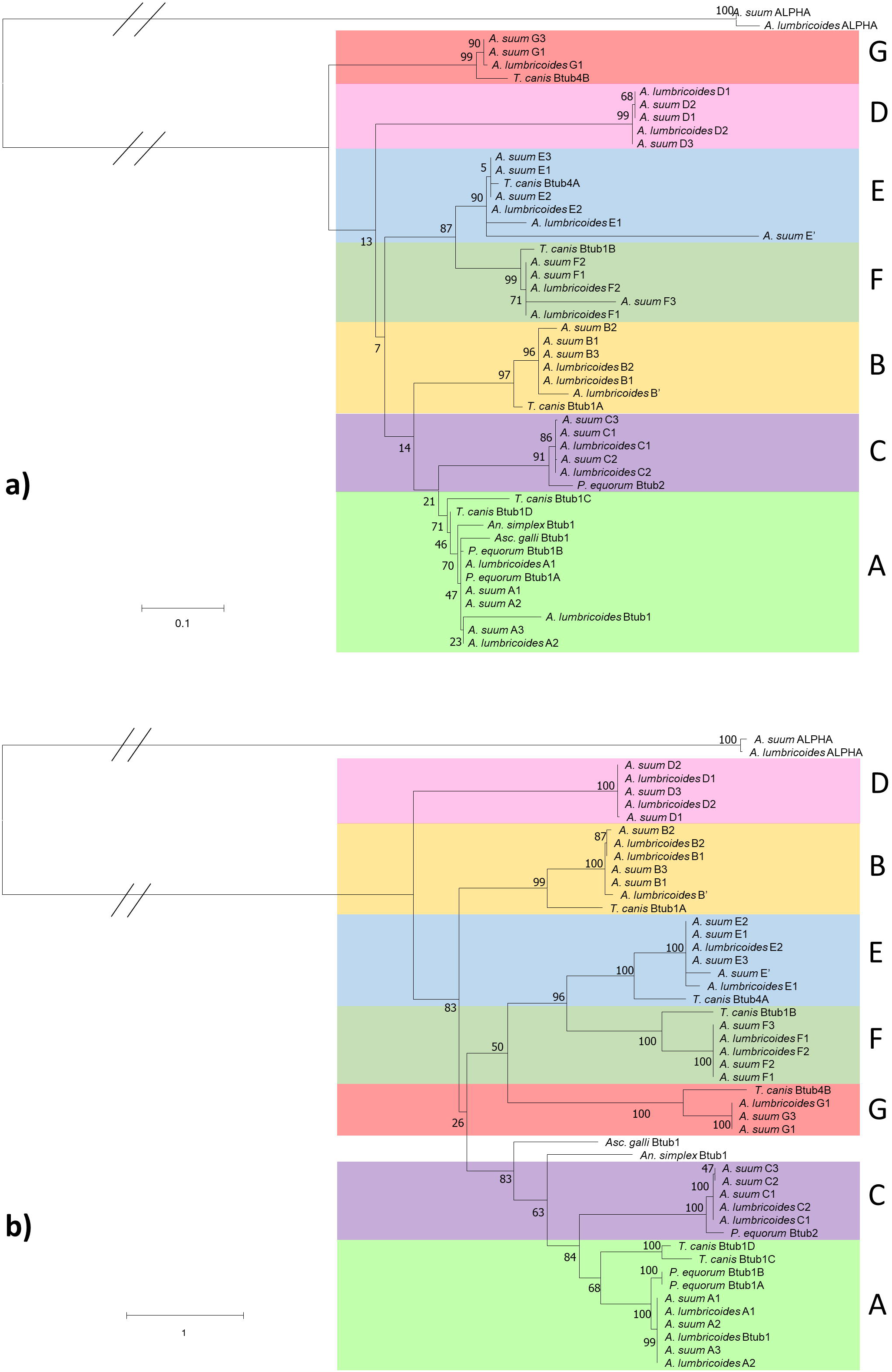
Phylogenetic reconstruction of Ascaridomorpha β-tubulins. Phylogenies show the relationship between each isotype from the *Ascaris* genomes as well as previously published Ascaridomorpha species β-tubulins. (a) shows the phylogeny reconstructed using the peptide sequences under the assumptions of the JTT+G model. (b) shows the nucleotide phylogeny reconstructed under the K2+G+I model. Both phylogenies underwent 1000 bootstraps. Bootstrap values are shown at each node. Sequences collected for members of the Ascaridomorpha have retained the nomenclature given in the database (e.g., *Toxocara canis* β-tub4B). The species included were *Anisakis simplex, Ascaridia galli, Parascaris equorum* and *Toxocara canis*. For the *Ascaris* sequences identified from the genomes each sample is named by species, isotype and then genome number (e.g., *Ascaris suum* C3 is the isotype C sequence from *A. suum* genome 3).

Only the seven isotypes that had homologues in both *Ascaris* species were used in further analysis. Isotype A clustered with sequences from other species, such as *Parascaris*, that have been previously linked with BZ interaction through gene expression studies, and is the isotype which is currently used in diagnostic tests for BZ resistance in *Ascaris* ^15,16,22,36,37^. For this reason, isotype A was used as the focus of molecular docking simulations. Interestingly the isotype previously designated as isotype-1 in *A. suum* did not fall within isotype A, but instead was found to be isotype C, suggesting the past labelling of this sequence as isotype-1 was incorrect ^20^.

### *In silico* docking shows similar binding for all β-tubulin isotypes

*In silico* ligand docking simulations were performed on the seven β-tubulin isotypes shared by both *Ascaris* species. An alignment of each isotype highlighting some active site amino acids can be seen in Figure 2. Five BZ drugs were docked into the active sites of each isotype and simulations showed a consistent trend between species, drug and isotype. However, the 3D structures and the 2D maps were not always in complete agreement when labelling hydrogen bonds (H-bonds). Hydrogen bond formation between BZs and amino acids Q134, E198 and V236 were the most common interactions and were consistently seen in all isotypes. Amino acid A315 and the amino acids at position 165 were also seen numerous times in docking poses. In the majority of cases amino acid 165 was a serine (S), although in isotypes E and F, amino acid 165 was asparagine (N) and threonine (T) respectively. These changes did not cause a change to the overall amino acid properties as all three amino acids are polar (neutral) hydrophobic amino acids, and similar interaction were observed between the drugs and all three amino acids. Other amino acids interacted with the BZs in some isotypes, and although these were not consistently seen, they could be of some importance and would require further investigation (Fig. 3).

**Figure 2:**
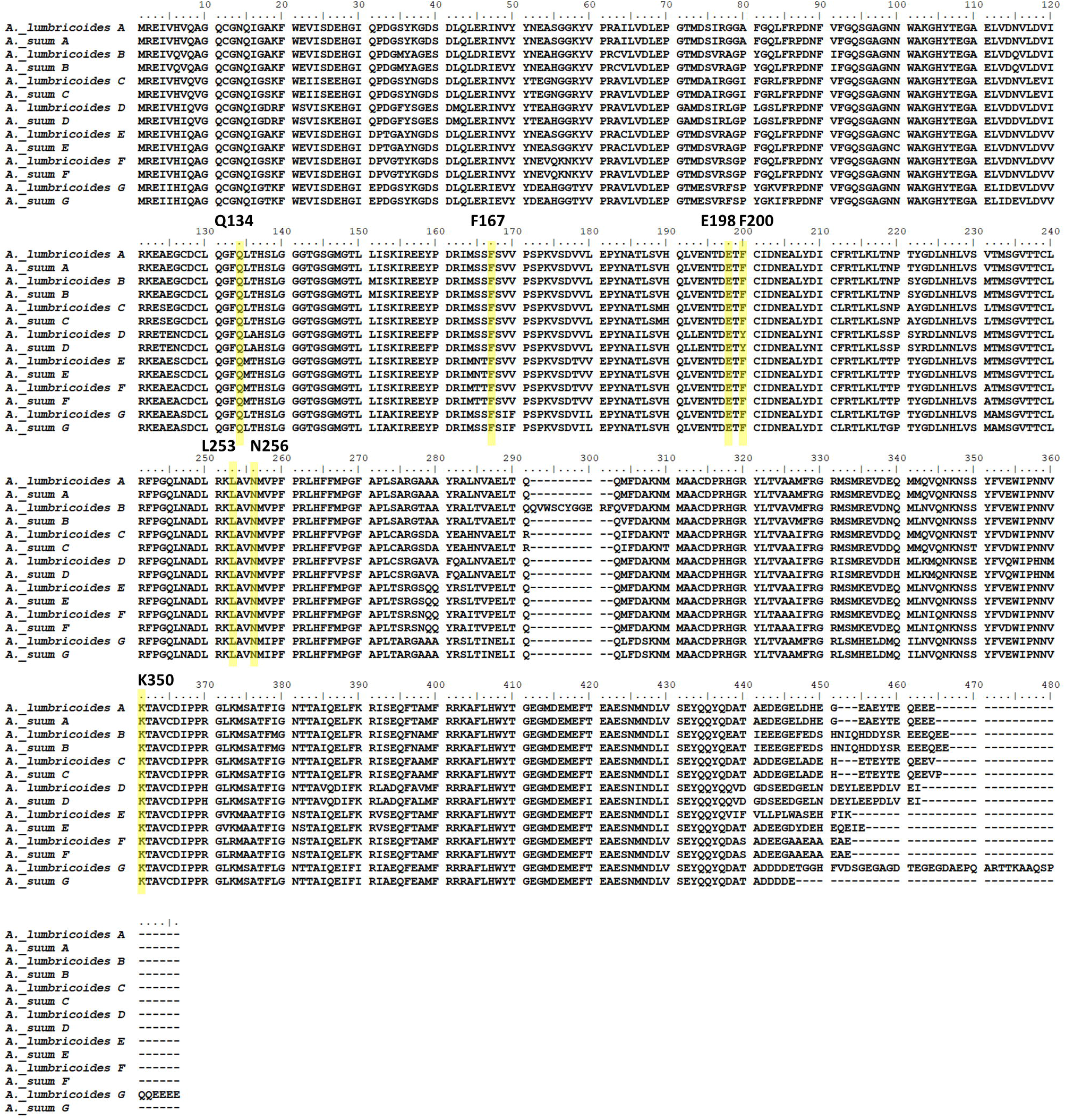
Representative amino acid alignment of *Ascaris* β-tubulins. Alignments for each *Ascaris* β-tubulin isotype used in docking simulations. The common resistance associated amino acids (F167, E198 and F200Y) and the amino acids that were found to interact with BZs (Q134, L253, N256 and K350) and may be of some importance are highlighted in yellow.

**Figure 3:**
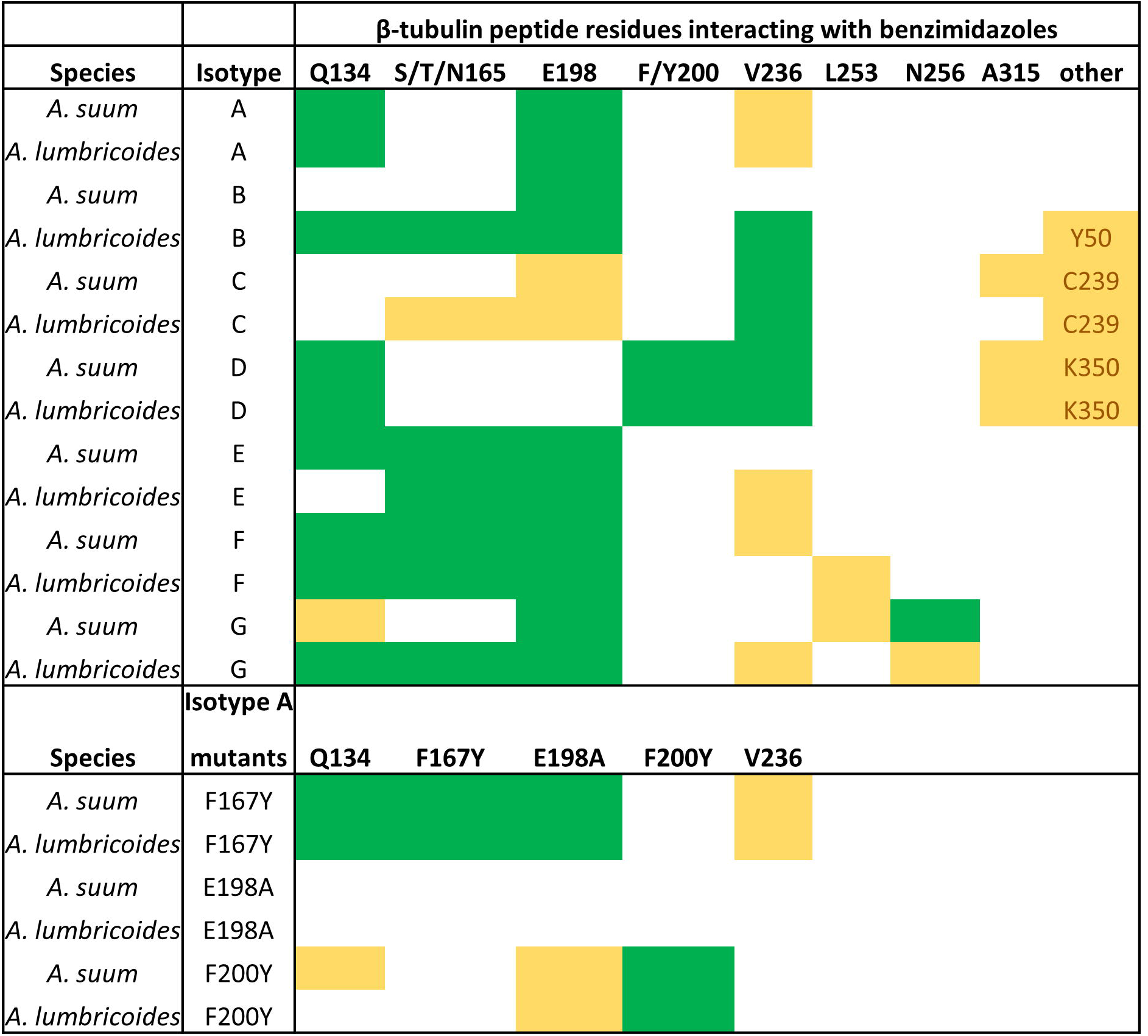
Ligand docking amino acid binding frequencies. The amino acids in *Ascaris lumbricoides* and *Ascaris suum* that form bonds with benzimidazole drugs in the ligand docking simulations are shown. Green represents an amino acid that interacts with more than one drug and yellow represents an interaction seen only once.

Isotype D had tyrosine (Y) at position 200 and this formed bonds with glutamate (E) at position 198. Isotype D was the only isotype to naturally contain tyrosine at position 200 which has been linked to resistance when seen in other β-tubulins ^21,27^. Recent work in *Parascaris* has shown that having tyrosine as the wildtype amino acid in this β-tubulin isotype is not restricted to *Ascaris* only ^38^.

It was only in isotype D and the mutated F200Y models that binding was seen between the drugs and amino acid 200. In the mutated F167Y protein models, the mutation of phenylalanine (F) to tyrosine resulted in extra bonds being formed with the drugs in most cases. In the mutated E198A models no bonds were formed with E198A in any drug model. The models with the F200Y mutation showed a bond between the mutated F200Y amino acid and E198. The full details of the binding of each drug to each individual isotype are provided in Supplementary Figures S2 – S21.

### Molecular dynamics simulations highlight BZ resistance mechanisms

Molecular dynamics simulations calculate the pressure and heat energies that are likely found within a physiological system and apply these to the protein-drug structure to mimic natural systems over a period of time to find the optimum binding poses. These simulations show how protein-drug interactions fluctuate over a period of time and give an indication of how these molecules may react in a physiological system. As the molecular docking simulations showed no difference between species or isotype, molecular dynamics simulations were performed only on *A. suum* isotype A. Simulations showed no major changes from the initial ligand docking. For *A. suum* isotype A, bonds between the protein and drugs formed with E198 in all models. Several other bonds were seen depending on the drug, but all models had similar binding affinities (Fig. 4, Table 1 and Table 2).

**Figure 4:**
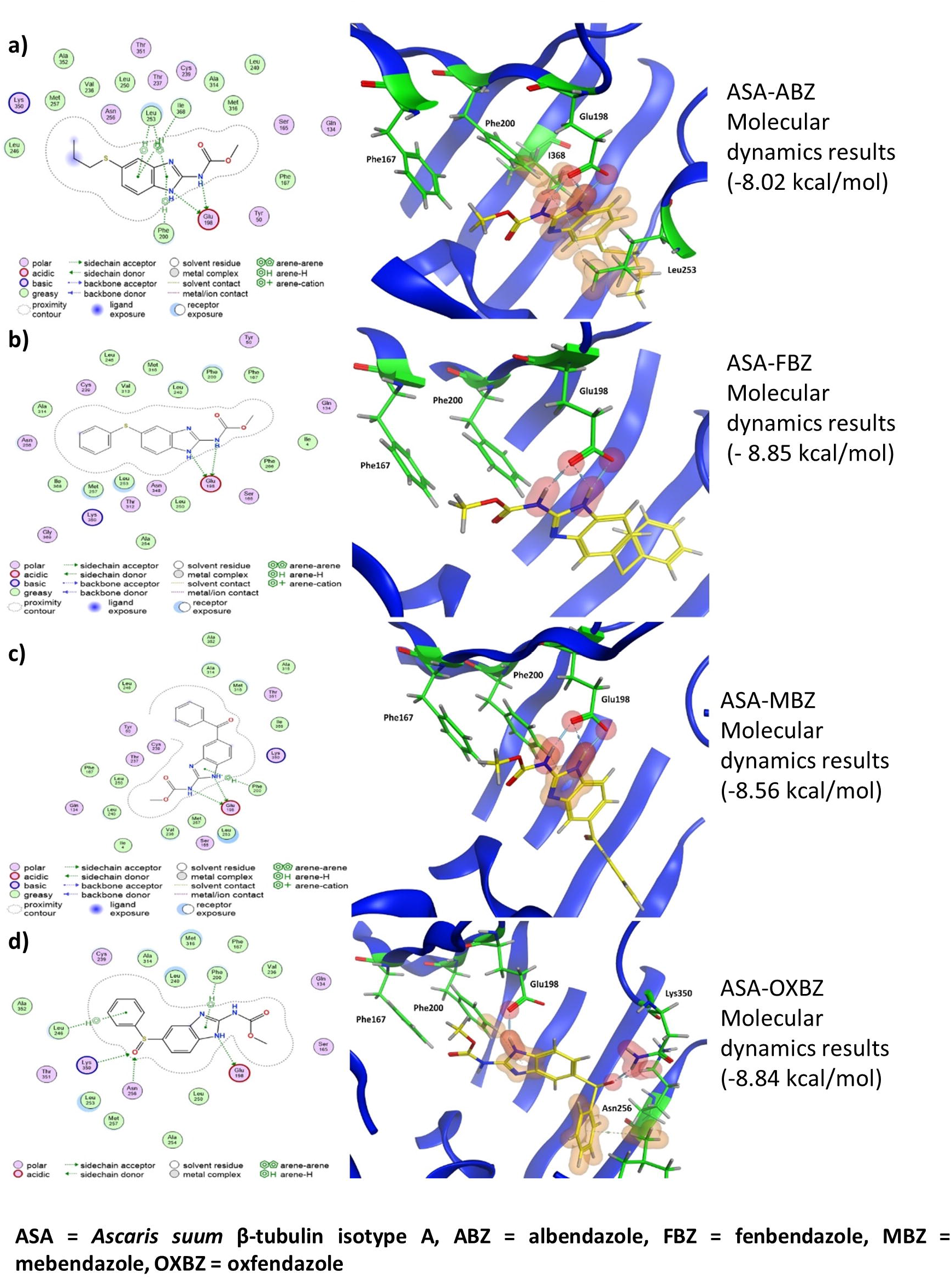
2D and 3D representations of the molecular dynamics simulations of *Ascaris suum* isotype A models with various benzimidazole drugs. The figure shows the protein-ligand interactions made in each model. In 2D models (left) bonds formed with amino acids are depicted with dashed lines with the specific type of bond indicated in the key. All amino acids shown in 2D models without bonds are predicted to interact via Van der Waals forces. In the 3D models (right) the protein structure is shown in ribbon format (blue) with only binding amino acids or the resistance associated amino acid shown in full (green). Hydrogen bonds (H-bonds) between protein and ligand are highlighted in red and arene bonds are highlighted in amber. The benzimidazole drugs are shown in yellow. Binding affinity is shown to the right of each model.

**Table 1:** Binding affinities of wildtype and mutated *Ascaris suum* and *Haemonchus contortus* β-tubulin proteins with benzimidazole drugs. Binding affinities of the protein-drug interactions from molecular dynamics simulations are measured in kcal/mol. The proteins used in these analyses were ASA, the three mutated ASA proteins, Hcon1 and the three mutated Hcon1 proteins. ASA = *Ascaris suum* β-tubulin isotype A, Hcon1 = *Haemonchus contortus* β-tubulin isotype-1, ABZ = albendazole, ABZSO = albendazole sulfoxide, FBZ = fenbendazole, MBZ = mebendazole, OXBZ = oxfendazole.

| Protein-drug model | Affinity (kcal/mol) |
| --- | --- |
| ASA-ABZSO | -8.16 |
| ASA-F167Y-ABZSO | -8.28 |
| ASA-E198A-ABZSO | -7.84 |
| ASA-F200Y-ABZSO | -8.07 |
| ASA-ABZ | -8.02 |
| ASA-FBZ | -8.85 |
| ASA-MBZ | -8.56 |
| ASA-OXBZ | -8.84 |
| Hcon1-ABZSO | -8.54 |
| Hcon1-F167Y-ABZSO | -8.22 |
| Hcon1-E198A-ABZSO | -7.12 |
| Hcon1-F200Y-ABZSO | -8.16 |

**Table 2:** Interactions between benzimidazole drugs and specific amino acid amino acids in Ascaris suum and Haemonchus contortus β-tubulin proteins. The proteins used in these analyses were *Ascaris suum* β-tubulin ASA, the three mutated ASA proteins, Hcon1 and the three mutated Hcon1 proteins. The drugs used were ABZ, ABZSO, FBZ, MBZ and OXBZ). The table shows the type of bonds formed (H – hydrogen bond, A – arene bond), the amino acid the bond is formed with and the drug used. The energy of the bonds between the amino acid and drug are given in kcal/mol, the distance between the bonded atoms is given in Angstroms (Å) and the number of bonds formed between the amino acid and the drug is shown (frequency). ASA = *Ascaris suum* β-tubulin isotype A, Hcon1 = *Haemonchus contortus* β-tubulin isotype-1, ABZ = albendazole, ABZSO = albendazole sulfoxide, FBZ = fenbendazole, MBZ = mebendazole, OXBZ = oxfendazole.

| Protein-drug model | Type | Amino acid | Drug | Energy | Distance (Å) | Frequency |
| --- | --- | --- | --- | --- | --- | --- |
| ASA-ABZSO | H | Glu198 | ABZSO | -9.4 | 3.06 | 2 |
|  | A | Phe200 | ABZSO | -0.5 | 4.65 | 1 |
|  | H | Val236 | ABZSO | -6.7 | 2.93 | 1 |
|  | A | Leu253 | ABZSO | -2.6 | 3.97 | 3 |
|  | H | Lys350 | ABZSO | -15.7 | 2.81 | 1 |
| ASA-F167Y-ABZSO | H | Glu198 | ABZSO | -15.9 | 2.76 | 2 |
|  | A | Phe200 | ABZSO | -1.2 | 4.13 | 2 |
|  | H | Met257 | ABZSO | -0.5 | 3.68 | 1 |
|  | H | Met316 | ABZSO | -1.4 | 3.94 | 2 |
| ASA-E198A-ABZSO | H | Gln134 | ABZSO | -5.3 | 2.86 | 1 |
|  | H | Leu253 | ABZSO | -1.3 | 3.03 | 1 |
| ASA-F200Y-ABZSO | H | Tyr50 | ABZSO | -2 | 2.77 | 1 |
|  | H | Glu198 | ABZSO | -18.4 | 2.77 | 2 |
| ASA-ABZ | H | Glu198 | ABZ | -18.3 | 2.82 | 3 |
|  | A | Phe200 | ABZ | -0.6 | 4.03 | 1 |
|  | A | Leu253 | ABZ | -1.5 | 3.68 | 2 |
|  | A | Ile368 | ABZ | -0.5 | 3.66 | 1 |
| ASA-FBZ | H | Glu198 | FBZ | -16.3 | 2.84 | 3 |
| ASA-MBZ | H | Glu198 | MBZ | -17.2 | 2.87 | 3 |
|  | A | Phe200 | MBZ | -0.5 | 3.67 | 1 |
| ASA-OXBZ | H | Glu198 | OXBZ | -7.6 | 2.89 | 1 |
|  | A | Phe200 | OXBZ | -0.5 | 4.2 | 1 |
|  | A | Leu246 | OXBZ | -0.8 | 3.99 | 1 |
|  | H | Asn256 | OXBZ | -6 | 2.82 | 1 |
|  | H | Lys350 | OXBZ | -1 | 3.52 | 1 |
| Hcon1-ABZSO | H | Glu198 | ABZSO | -16.9 | 2.87 | 3 |
| Hcon1-F167Y-ABZSO | H | Glu198 | ABZSO | -11.6 | 2.75 | 2 |
|  | A | Leu253 | ABZSO | -0.7 | 3.99 | 1 |
|  | H | Asn256 | ABZSO | -5.6 | 2.82 | 1 |
| Hcon1-E198A-ABZSO | H | Leu253 | ABZSO | -3.5 | 2.85 | 1 |
| Hcon1-F200Y-ABZSO | H | Glu198 | ABZSO | -0.5 | 3.56 | 1 |
|  | H | Cys239 | ABZSO | -3.1 | 2.98 | 1 |
|  | H | Asn256 | ABZSO | -7.6 | 2.92 | 2 |
|  | H | Met316 | ABZSO | -2 | 3.54 | 2 |

In the E198A mutation model there was a reduced binding affinity and complete loss of bonding with amino acid E198A, although weaker bonds still formed with other common amino acids (Fig. 5c, Table 1 and Table 2). No difference in drug interactions were seen in the F200Y model compared to the wildtype model, although the mutated F200Y amino acid did form a self-binding interaction with E198 (Fig. 5e). In the F167Y model the additional bond formed with F167Y in the ligand docking models was not seen and there was no direct effect of this mutation on drug binding (Fig. 5).

**Figure 5:**
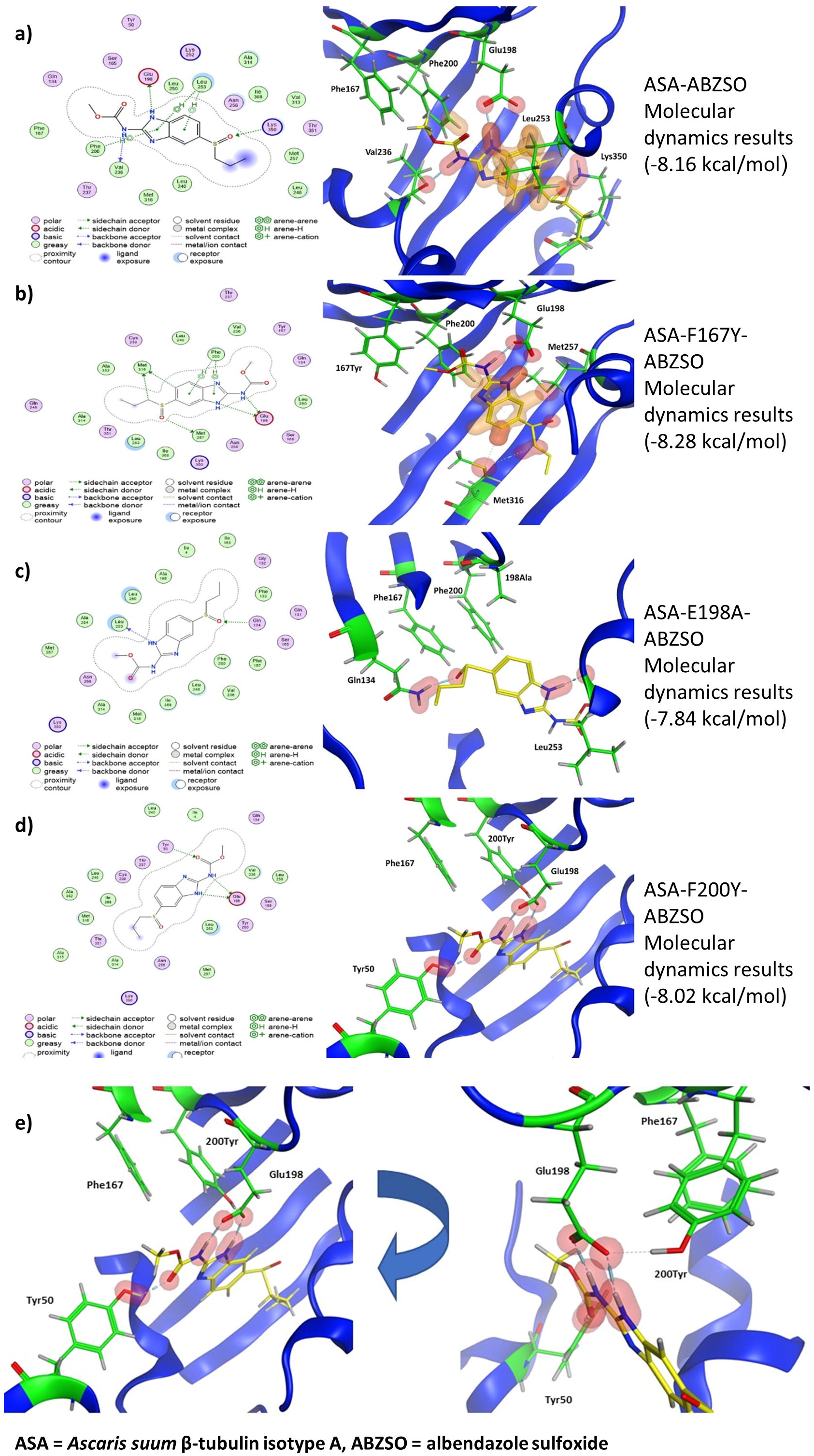
2D and 3D representations of the molecular dynamics simulations of *Ascaris suum* β-tubulin isotype A wildtype and mutant models with albendazole sulfoxide. The figure shows the protein-ligand interactions made within each model. In 2D models (left) bonds formed with amino acids are depicted with dashed lines with the specific type of bond indicated in the key. All amino acids shown in 2D models without bonds are predicted to interact via Van der Waals forces. In the 3D models (right) the protein structure is shown in ribbon format (blue) with only binding amino acids or the resistance associated amino acid shown in full (green). Hydrogen bonds (H-bonds) between protein and ligand are highlighted in red and arene bonds are highlighted in amber. The drug ABZSO is shown in yellow. Binding affinity is shown to the right of each model. Models shown are (a) wildtype ASA, (b) mutated 167Y ASA, (c) mutated 198A ASA and (d) mutated 200Y ASA. (e) shows the H-bonds between the drug and Y50 and E198 as seen in (d) but also shows a rotated view of this model so that the bond between E198 and 200Y is made visible.

Resistance to BZs has been best documented in *H. contortus* and previous *in silico* modelling simulations have been performed on this species to explore BZ resistance mechanisms ^32,34^. For these reasons we performed molecular dynamics simulations on *H. contortus* β-tubulin isotype-1 to compare our results with previous studies in this model organism. All simulations using *H. contortus* models compared well with *A. suum* models (Fig. 6). For the wildtype susceptible protein, H-bonds formed with E198 (Fig. 6a). There was no direct interaction observed between the drug and the F167Y amino acid amino acid (Fig. 6). In the E198A models reduced binding affinity was observed, and a loss of interaction with E198A with only weak bonds formed with other amino acids (Fig. 6c, Table 2). Finally, the F200Y mutation resulted in interactions between E198 and the F200Y amino acid amino acids and drug interactions with E198 were weakened (Fig. 6d, Table 2).

**Figure 6:**
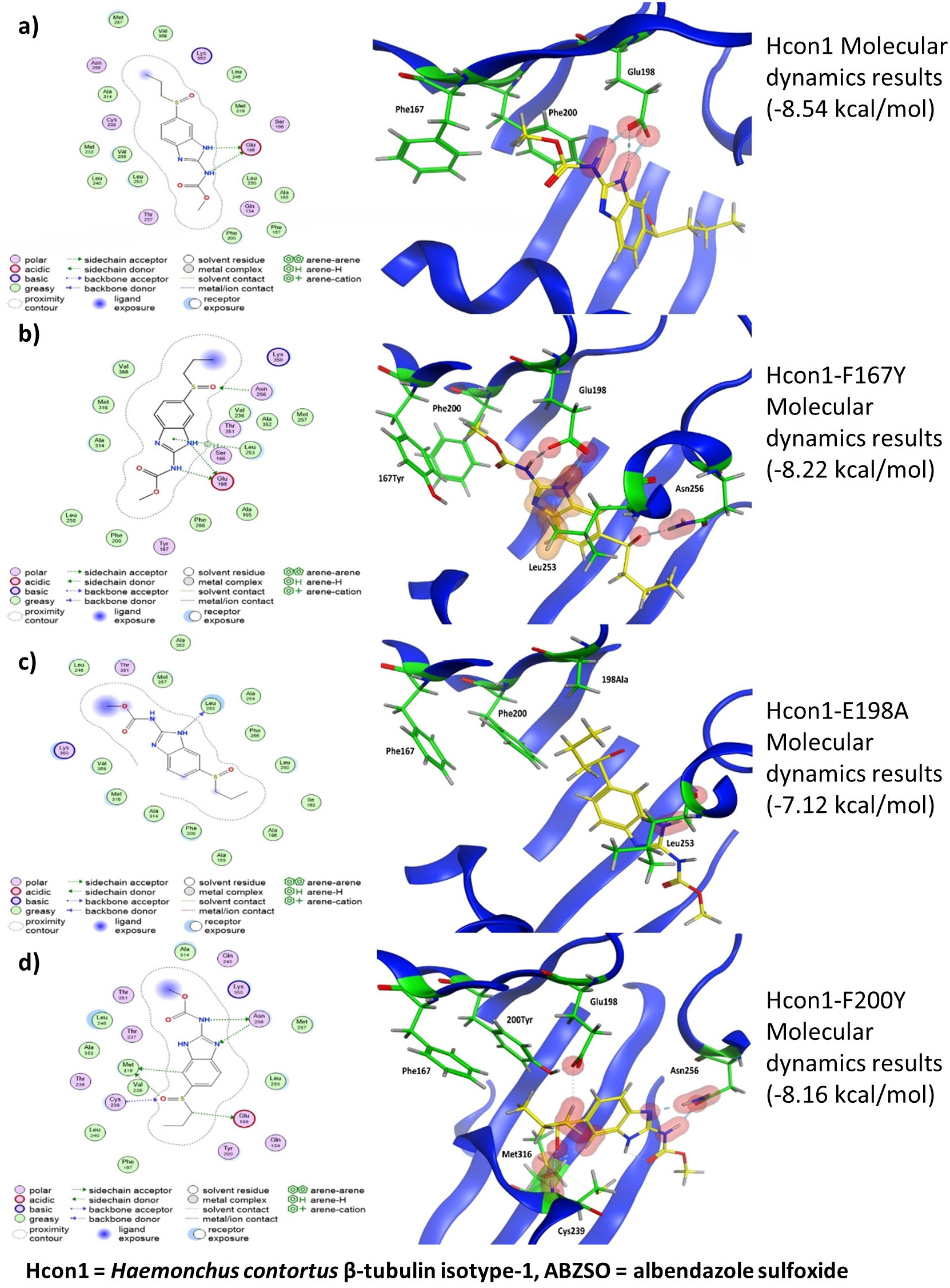
2D and 3D representations of the molecular dynamics simulations of *Haemonchus contortus* β-tubulin isotype 1 wildtype and mutant models with albendazole sulfoxide. The figure shows the protein-ligand interaction made in each model. In 2D models (left) bonds formed with amino acids are depicted with dashed lines with the specific type of bond indicated in the key. All amino acids shown in 2D models without bonds are predicted to interact via Van der Waals forces. In the 3D models (right) the protein structure is shown in ribbon format (blue) with only binding amino acids or the resistance associated amino acid shown in full (green). Hydrogen bonds (H-bonds) between protein and ligand are highlighted in red and arene bonds are highlighted in amber. The drug albendazole sulfoxide (ABZSO) is shown in yellow. Binding affinity shown to the right of each model. Models shown are (a) wildtype Hcon1, (b) mutated 167Y Hcon1, (c) mutated 198A Hcon1 and (d) mutated 200Y Hcon1.

## Discussion

The widespread resistance to BZs in ruminant nematodes such as *H. contortus* has illustrated the effects that resistance can have on both animal health and economic returns ^39^. We have not yet seen widespread resistance to BZs in *Ascaris i*n either humans or pigs, although, with increasing drug pressure to reach the 2030 World Health Organisation targets, limited studies on drug efficacy in either humans or pigs, and limited alternative treatments, a better understanding of the mechanisms leading towards BZ resistance in *Ascaris* is urgently required. This work identified seven β-tubulin isotypes shared by both *Ascaris* species considered here, and compared, *in silico*, BZ interactions between them. We observed that all β-tubulin isotypes are predicted to interact with BZs in a similar manner, except for one isotype that contains a resistance-associated amino acid at position 200 in its wildtype protein. *In silico* ligand docking and molecular dynamics simulations highlighted E198 as a key amino acid in BZ-binding, with E198A mutations leading to weaker protein-drug interaction. We also found that the common resistance associated F200Y mutation acts indirectly by binding to E198 and reducing drug stability within the binding pocket.

By utilising multiple databases, we were able to identify seven β-tubulin isotypes from both *A. suum* and *A. lumbricoides*. Phylogenetic analysis showed that *Ascaris* β-tubulin isotypes were shared with other Ascaridomorpha species, and it is isotype A that is used as a marker of BZ resistance and is usually referred to as isotype-1 ^15,16,22,36,37^. The identification of isotype A as the main group involved in BZ interaction allowed *in silico* work to focus on this isotype. Concurrent work in *Ascaris* by Roose *et al*. ^40^ also found these same β-tubulin isotypes in both species and identified isotype A as the isotype used in previous surveillance studies. Isotype A was shown to be the most highly expressed β-tubulin isotype and therefore one of the main isotypes likely to be involved in BZ interaction ^40^. Whilst the expression levels of the β-tubulin isotypes in *Ascaris* are now known, the contribution of these to drug mechanisms of action have not yet been defined ^40^. Our work has shown that the drug interaction with these isotypes does not differ on the whole, with the exception of isotype D. Therefore, it likely that the contribution of each isotype to drug-binding is relative to the expression level during the different stages of the *Ascaris* life-cycle.

The most common binding amino acids predicted from molecular dynamics simulations were E198, L253 and N256.Several other amino acids interacted with the BZs, although these were not consistent. Most of these interactions had weak binding affinity, although N256 and K350 were shown to form stronger bonds and could be of potential importance. It has been assumed that E198 is the key binding amino acid for BZs, and indeed the key role E198 has in BZ binding and the self-binding interaction between amino acid E198 and F200Y in the mutated models was observed ^33,34^. By investigating binding energies at each amino acid it has been shown here that the bonds between the BZs and E198 are much weaker in the *H. contortus* F200Y mutated models than in *Ascaris* models. This has not been demonstrated before, and adds further evidence to the theory that interactions between E198 and F200Y destabilize BZ binding ^34^.

Our results suggest that E198 is the key amino acid in β-tubulin for BZ binding in *Ascaris*, as interactions were seen in every model except for the mutated E198A structure. Bonds with E198 also showed the strongest binding affinity; at least three times as strong as any other amino acid interaction in most cases. In models that contained the E198A mutation, the change led to a loss of interaction at this important site. In F200Y simulations, the self-binding between E198 and F200Y was observed, which could lead to the blocking or destabilising of interactions between BZs and E198, resulting in resistance to BZs. Interestingly, the binding energy between BZ and E198 in the *A. suum* F200Y models was not reduced as much as it was for *H. contortus*. In F167Y models there was no clear change, and this lack of any clear negative effect may explain why the F167Y mutation has been found in field isolates of *A. lumbricoides* without any effect on drug susceptibility ^23^. However, in *H. contortus* F167Y models, there was also no effect on binding, although in *H. contortus* this mutation is known to cause resistance, which suggests that the models may still be unable to predict more complicated mechanisms of resistance. It has been hypothesised that the F167Y mutation leads to self-binding with amino acids that close off the binding pocket and prevent the drugs from entering ^34^. In our work no such self-binding could be seen between the tyrosine at position 167 and any other amino acids.

Benzimidazole resistance is common for *H. contortus* and other clade V nematodes but is yet to become a common problem for *Ascaris*. In all the searches for drug resistance in *Ascaris* to date only the three common resistance associated mutations, F167Y, E198A and F200Y have been investigated, which means the contributions of other mutations that may affect the BZ susceptibility will be missed. There are reports of Ascarid helminths displaying reduced susceptibility to BZ, but do not contain these classical mutations, and hence there is a possibility that there may be other mechanisms or mutations involved in BZ resistance ^15,38^. In this study several other amino acids were identified as possible candidates, such as N256 and K350 (see Fig. 3 and Table 2 for full list of interacting amino acids), that may play an important role in drug binding and may lead to BZ resistance if mutations occur.

In conclusion, we have identified the full repertoire of β-tubulin genes from *A. lumbricoides* and *A. suum* and have shown that whilst almost all have the potential to interact with BZs, there is one isotype, isotype A, that is likely key to BZ binding. By identifying the importance of isotype A, our findings will allow future studies to refine and focus their approach to studying the effects of BZs in non-clade V nematodes and monitor resistance development. Our results show that E198 is a vital amino acid for BZ binding of β-tubulins in *Ascaris*, as has been seen for other helminths species; and the E198A and F200Y mutations both take effect by disrupting this key anchor point. However, it appears that in *H. contortus* the F200Y mutation causes more disruption to E198 binding than is seen in *Ascaris* and could be the key difference between the two groups of parasites. This new information may prove to be of significance for the molecular monitoring and modelling of resistance in *Ascaris* and could be key to understanding why resistance is so commonly reported in strongyle nematodes but as yet rarely so in Ascaridomorpha.

## Methods

Five *Ascaris* genomes: two for *A. lumbricoides* (GCA_000951055.1 and GCA_015227635.1) and three for *A. suum* (GCA_000298755.1, GCA_000187025.3 and GCA_013433145.1) were analysed to identify potential β-tubulin isotypes. Based on previous literature it was found that one *A. lumbricoides* β-tubulin gene had been characterised and deposited in the National Center for Biotechnology Information (NCBI) along with 21 β-tubulin sequences from *A. suum* ^20,22,41^. These sequences were retrieved from the database and the *A. suum* sequences were aligned with each other to remove the partial sequences that were duplicates of the longer sequences. The β-tubulin gene from *A. lumbricoides* (EU814697.1) retrieved from NCBI was used to carry out BLAST ^42^ searches against the three available *Ascaris* genomes in WormBase-Parasite (GCA_000187025.3, GCA_000298755.1 and GCA_000951055.1) ^43,44^. To ensure that no β-tubulin genes had been missed, the paralogues of each gene were checked, and the search term “tubulin beta” was used for each annotated genome. An α-tubulin sequence for each species was also retrieved to be used as the outgroup in further analysis.

Exonerate v2.2.0 ^45^ protein2genome was used to identify any β-tubulin genes within the *Ascaris* genomes that had not been detected by BLAST or in the genome annotation. Each isotype retrieved from the database search was run against all three genomes with the best 10 results being saved from each test. This number of tests were saved as we found up to eight potential isotypes from the database searches, and this allowed for the potential of at least two further sequences to be identified. Any new sequence found by Exonerate was tested in the Conserved Domain Database ^46^ to check that the sequence was a β-tubulin gene and then any new sequences predicted to be β-tubulins were added to the β-tubulin dataset. The two newest genomes (*A. lumbricoides* GCA_015227635.1 and *A. suum* GCA_013433145.1) had not been fully annotated and so Exonerate protein2genome was used to identify β-tubulin genes.

After all the sequences, both nucleotide and peptide, had been collected, they were aligned with tubulin sequences from other Ascaridomorpha using the MUSCLE server (available at: https://www.ebi.ac.uk/Tools/msa/muscle/ [Accessed 09 December 2020]) ^47^ and a maximum likelihood phylogeny was created with MEGA version X ^48^, using the JTT+G model for the amino acid sequences and the K2+G+I model for the nucleotide sequences. Each phylogeny was bootstrapped 1000 times. Genomes are numbered in the phylogenies as follows: *A. lumbricoides* 1 (GCA_000951055.1); *A. lumbricoides* 2 (GCA_015227635.1); *A. suum* 1 (GCA_000298755.1); *A. suum* 2 (GCA_000187025.3) and *A. suum* 3 (GCA_013433145.1). The peptide sequences of these genes were used to create homology models.

### Homology models

Homology models were created for all β-tubulin isotypes of *A. lumbricoides* and *A. suum* using SWISS-MODEL server (available at: https://swissmodel.expasy.org/ [Accessed 22 February 2021]) ^49,50^. The β-tubulin crystal structure 6fkj was used as the reference structure. This structure was chosen as it is an experimentally determined crystal structure containing multiple α/β-tubulin dimers with a ligand bound in the colchicine binding site in a similar way as predicted previously for BZs. The ligand bound to this structure is a cyclohexanedione derivative called TUB075 used as a tubulin targeting, antiproliferation cancer drug ^51^. Sequences for β-tubulin isotype A were edited to provide sequences with the common BZ resistance associated mutations (F167Y, E198A and F200Y) ^17,24^. These were again submitted to SWISS-MODEL to create homology models. This isotype was used as it was this isotype that had been identified as being highly expressed in previous studies of Ascaridomorpha ^37^.

### Quality checks

Homology models were submitted to multiple servers for quality checks to confirm the validity of structures created in SWISS-MODEL and help predict any potentially erroneous sites. ProSA-web (Protein Structure analysis) server (available at: https://prosa.services.came.sbg.ac.at/prosa.php [Accessed 15 March 2021]) ^52,53^ assesses protein model quality. Verify3D ^54,55^ compares the 3D structure of the model to the 1D peptide sequence. PROCHECK v3.5 ^56^ analyses the structural geometry of the protein structures using Ramachandran plots. Both Verify3D and PROCHECK are part of the UCLA SAVES v6.0 server (available at: https://saves.mbi.ucla.edu/ [Accessed 15 March 2021]).

### Energy minimisation

Structures were minimised using the YASARA energy minimization server (available at: http://www.yasara.org/minimizationserver.htm [Accessed 25 February 2021]) ^57^. This server uses the YASARA forcefield to optimise the positions of atoms and reduce interatomic energies. After all structures were minimised quality checks were performed again. The quality checks of minimised homology models show acceptable results; with Z-scores within the expected range in ProSA, verify3D scores over 80% and no errors found with PROCHECK.

### *In-silico* ligand docking

3D ligand structure files for commonly used BZ drugs were downloaded from PubChem ^58^ in SDF format. The drugs used include three of the most commonly used BZ, albendazole (ABZ), mebendazole (MBZ) and fendazole (FBZ), as well as albendazole sulfoxide (ABZSO) and oxfendazole (OXBZ), which are the active metabolites of ABZ and FBZ respectively. These were converted into pdb format using Pymol v2.3.4 ^59^. Pdb structures of ligands were uploaded to Autodock tools v1.5.6 ^60,61^. The number of allowable rotatable bonds was set to maximum, and structures were saved in pdbqt format suitable for docking simulations. Protein models were uploaded to Autodock tools to be prepared for docking simulations. Water was deleted from the protein structures; polar hydrogens were added, and structures were saved in pdbqt format. The docking grid was centred on amino acid 200 of the protein as this is the primary amino acid believed to be associated with BZ resistance. Grid spacings were set to 1 Angstrom (Å) and box size was set to 24Å for x, y and z sizes. This grid box encased all three resistance associated amino acids within a small pocket of the protein and the co-ordinates of the box were saved for later use.

Autodock vina v1.1.2 ^62^ was used to perform *in-silico* ligand docking simulations between the β-tubulin isotypes and BZ drugs, using the grid co-ordinates and spacings to identify the target binding region and an exhaustiveness level of 8. Docking results were opened in Pymol to view the 3D structure and interactions. Polar contacts between the drug and proteins were identified and protein-ligand complexes were exported in pdb format. Protein-ligand complexes were opened in Discovery studio v20.1.0.19295 ^63^ to create 2D ligand interaction maps which show multiple types of interaction between the protein and ligand in a clear and easily read format.

### Molecular dynamics

Molecular dynamics simulation were carried out using Molecular Operating Environment (MOE) 2020.01 ^64^. The β-tubulin structures were optimised using the Protonate3D method with default settings in MOE. The site finder algorithm was then implemented to identify binding pockets within the protein. The pocket corresponding to the known binding region of BZs was selected and dummy atoms were inserted as markers for the docking. Initial docking simulations were run for each BZ with the *A. suum* β-tubulin isotype A using the dummy atoms as the site of binding. The initial scoring of docking poses used the London dG method to identify the best 30 ligand poses. This was followed by final scoring of the best 10 poses using GBVI/WSA dG method. Only ABZSO was used for the mutated versions of isotype A. The results of the MOE docking were then used for molecular dynamics simulations using the NPA algorithm and the Amber10: EHT forcefield using default configurations. Structures were equilibrated for 100 picoseconds (ps) at 300°K before a production run of 500 ps at 300°K with a time step of 0.002 ps. Once completed, the binding energy of each interacting amino acid and the overall energy in the binding pocket is calculated.

A selection of timesteps were taken every 50 ps. For each of the selected timesteps the ligand was constrained, and the structure was minimised to give the binding affinity of the ligand. The pose with the strongest binding affinity was then selected as the final result and 2D and 3D representations of the final model were saved. Due to the similarity between species and β-tubulin isotypes, only *A. suum* isotype A complexes, and their mutated forms were subject to this analysis. As a point of comparison with a better studied organism, *H. contortus* β-tubulin 1 (ACS29564.1), and mutated versions of this protein containing the BZ resistance associated SNPs were also analysed by molecular dynamics simulations.

## Data availability

The genomic datasets analysed during the current study are available in the Wormbase-Parasite and NCBI repositories, https://parasite.wormbase.org/Ascaris_lumbricoides_prjeb4950/Info/Index/; https://parasite.wormbase.org/Ascaris_suum_prjna62057/Info/Index/; https://parasite.wormbase.org/Ascaris_suum_prjna80881/Info/Index/; https://www.ncbi.nlm.nih.gov/genome/350?genome_assembly_id=925559; https://www.ncbi.nlm.nih.gov/genome/11969?genome_assembly_id=1482971.

## Acknowledgements

This study was funded by the Kenneth Longhurst legacy PhD studentship Award from the University of Surrey.

## Author contributions statement

B.P.J, M.B., A.H.M.V., E.J.L. designed the study. B.P.J. performed experiments. B.P.J, M.B., A.H.M.V. contributed to analysis of results. B.P.J, M.B., A.H.M.V., E.J.L contributed to writing manuscript. All authors have reviewed the final manuscript.

