## Supplementary figures and images for "Identification of key interactions of benzimidazole resistance-associated amino acid mutations in *Ascaris* β-tubulins by molecular docking simulations"

### Supplematary Figure S1

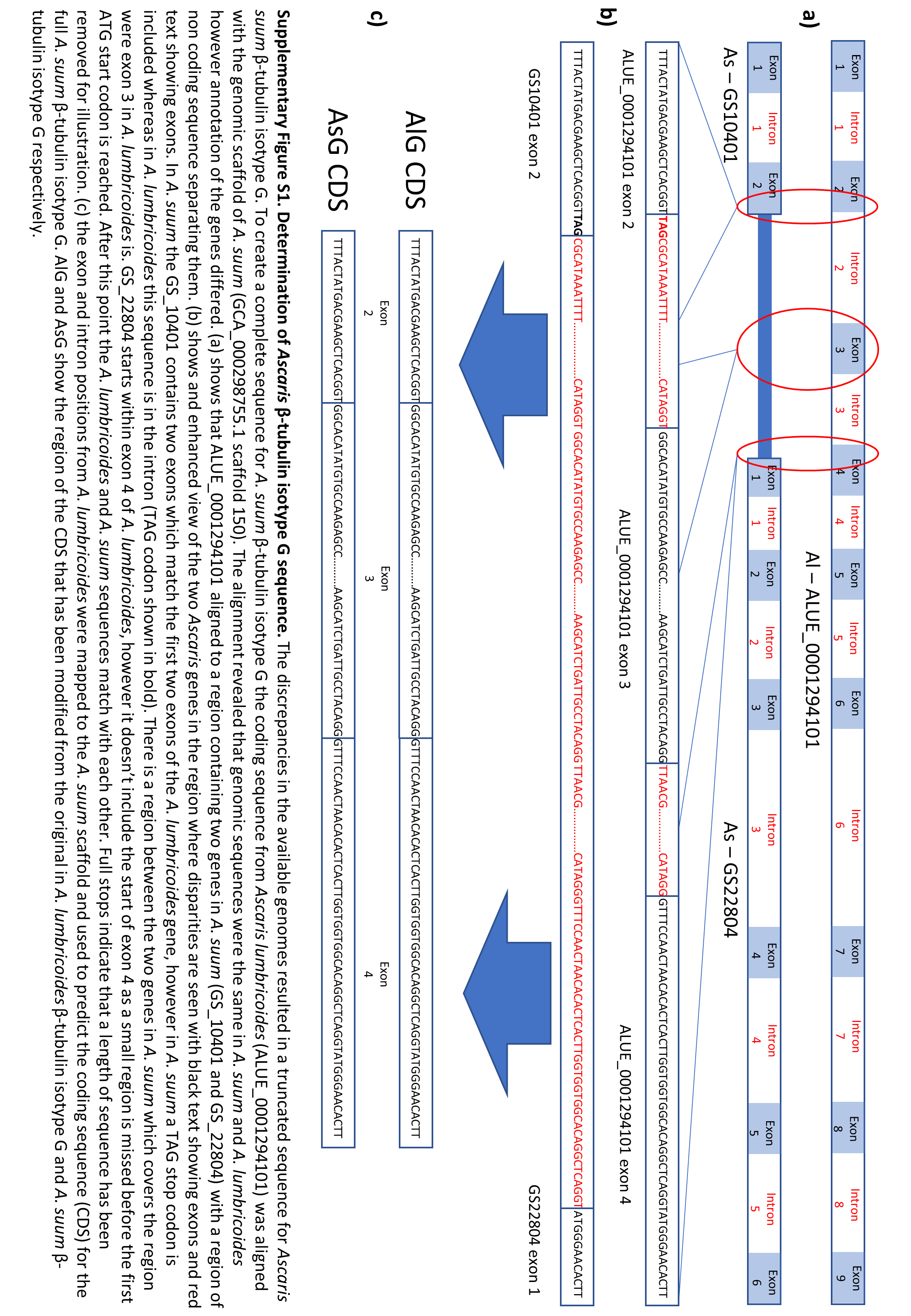
