## Supplementary Figures S2-21 for "Identification of key interactions of benzimidazole resistance-associated amino acid mutations in *Ascaris* β-tubulins by molecular docking simulations"

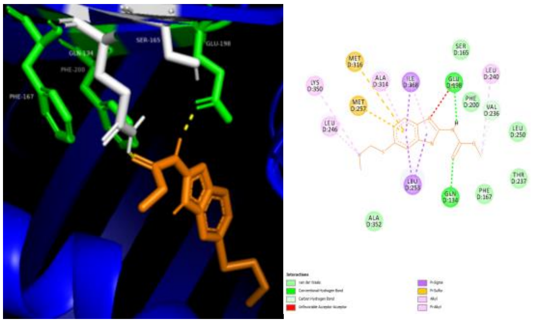

ASA-ABZ -5.8 kcal/mol

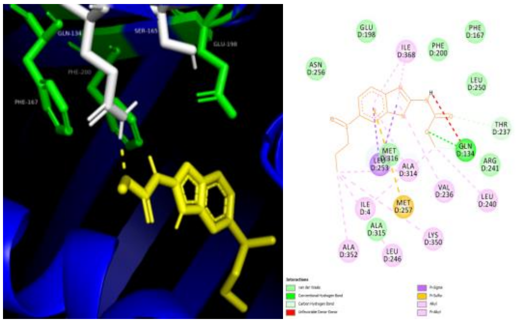

ASA-ABZSO -5.7 kcal/mol

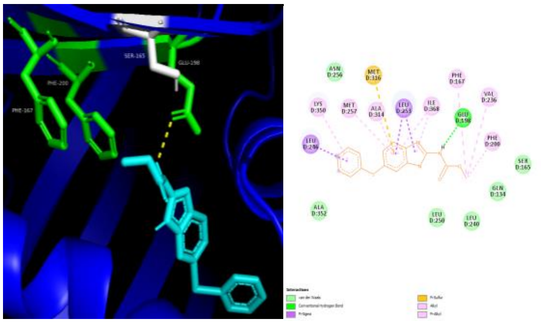

ASA-FBZ -6 kcal/mol

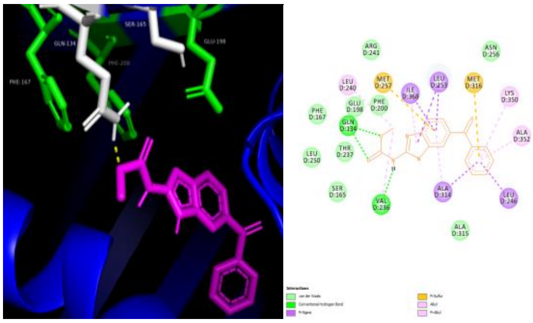

ASA-MBZ -6.2 kcal/mol

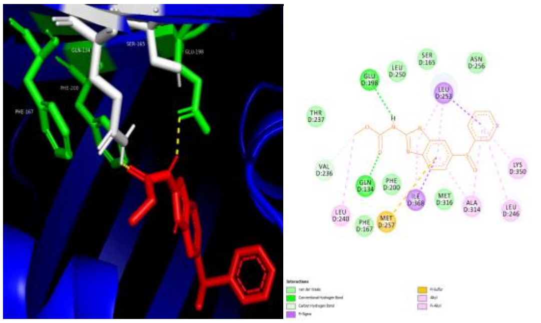

ASA-OXBZ -6.2 kcal/mol

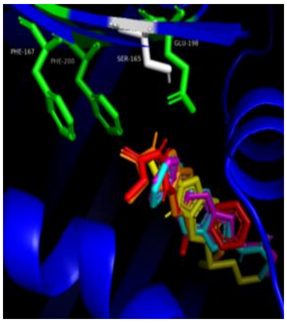

ASA-all drugs

**Supplementary Figure S2: Autodock vina docking results for *A. suum* isotype A and several benzimidazole drugs**. The drugs used are albendazole (ABZ), albendazole sulfoxide (ABZSO), fenbendazole (FBZ), mebendazole (MBZ) and oxfendazole (OXBZ). 3D and 2D models are shown for each docking result. Binding affinities are shown underneath each model. In ABZ docking H-bonds form with Q134 and E198 and an additional unfavourable acceptor-acceptor bond with E198 is also seen in the 2D models. For ABZSO the only H-bond seen is with Q134 in both models, and again an unfavourable bond is seen in the 2D model, this time a donor-donor bond is made with Q134. In both FBZ models the only bond found is with E198. For MBZ the 3D model shows a single H-bond with Q134 whereas in the 2D model two bonds are made with Q134 and one is made with V236. Finally, in both OXBZ models one H-bond is formed with Q134 and E198

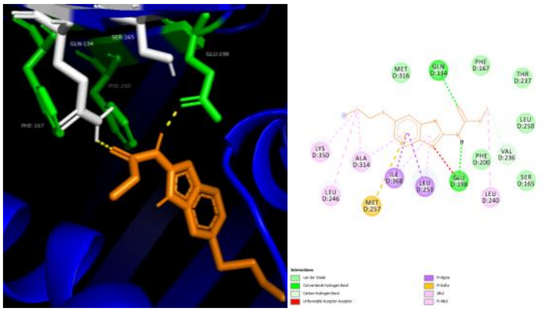

ALA- ABZ -5.7 kcal/mol

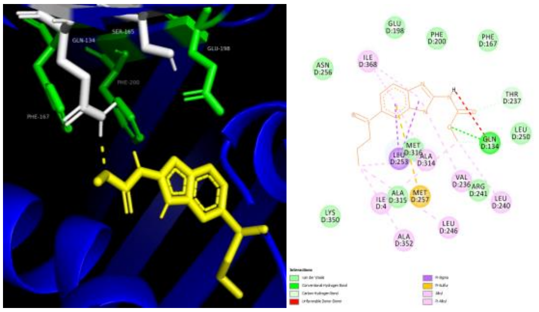

ALA- ABZSO -5.7 kcal/mol

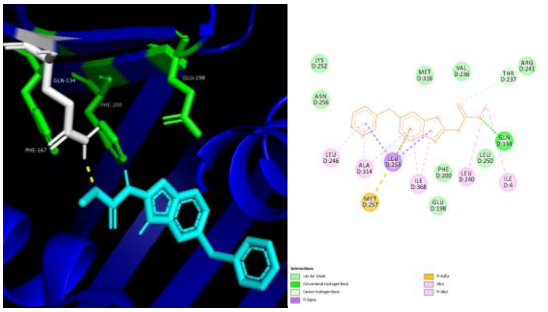

ALA- FBZ -6.1 kcal/mol

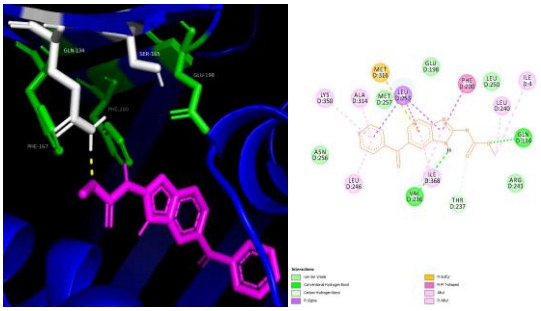

ALA- MBZ -5.8 kcal/mol

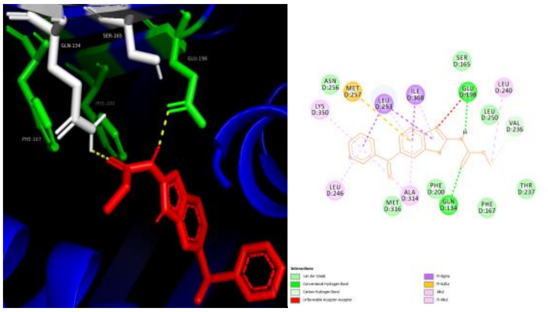

ALA- OXBZ -6.3 kcal/mol

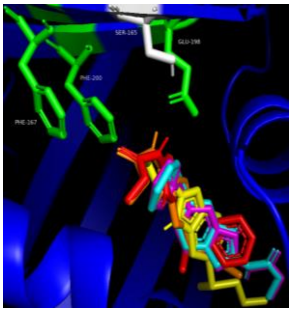

ALA- all drugs

**Supplementary Figure S3: Autodock vina docking results for *A. lumbricoides* isotype A and several benzimidazole drugs.** The drugs used are albendazole (ABZ), albendazole sulfoxide (ABZSO), fenbendazole (FBZ), mebendazole (MBZ) and oxfendazole (OXBZ). 3D and 2D models are shown for each docking result. Binding affinities are shown underneath each model. In ABZ docking H-bonds form with Q134 and E198 and an additional unfavourable acceptor-acceptor bond with E198 is also seen in the 2D models. For ABZSO the only H-bond seen is with Q134 in both models, and again an unfavourable bond is seen in the 2D model, this time a donor-donor bond is made with Q134. In the FBZ binding models single H-bonds are predicted with L253 and N256. For MBZ the 3D model shows a single H-bond with Q134 whereas the 2D model shows an extra bond is made with V236. Both models for OXBZ show one H-bond with Q134 and E198, yet in the 2D model and additional unfavourable acceptor-acceptor bond with E198.

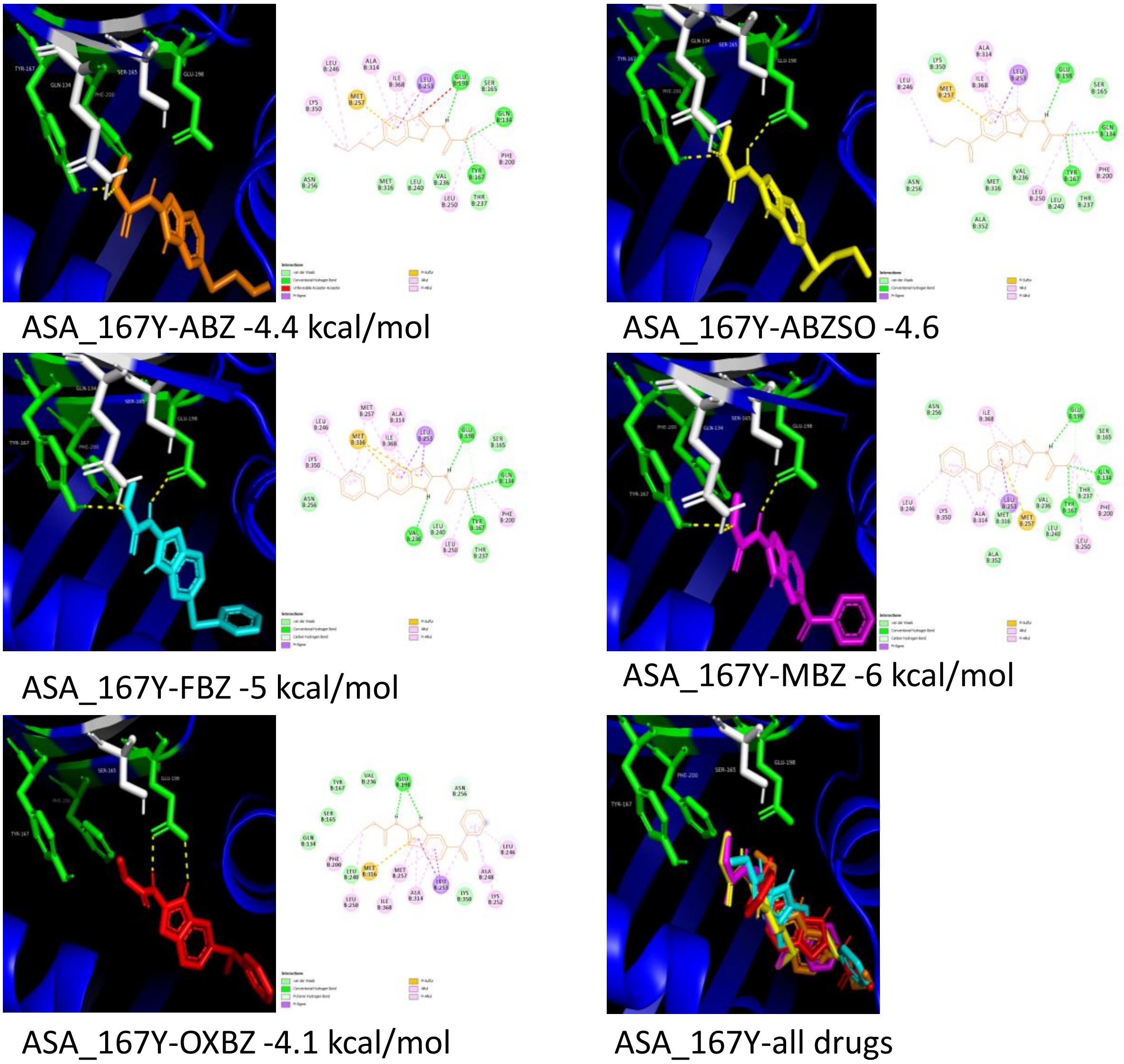


**Supplementary Figure S4: Autodock vina docking results for *A. suum* 167Y mutated isotype A and several benzimidazole drugs**. The drugs used are albendazole (ABZ), albendazole sulfoxide (ABZSO), fenbendazole (FBZ), mebendazole (MBZ) and oxfendazole (OXBZ). 3D and 2D models are shown for each docking result. Binding affinities are shown underneath each model. In these mutated 167Y models we see the larger tyrosine amino acid forming bonds with the drugs whereas the natural phenylalanine amino acid does not interact in any other amino acids. In ABZ 3D model we find H-bond formation with Q134 and 167Y. In the 2D model, however, we also see an additional H-bond and unfavourable acceptor-acceptor bond with E198. For ABZSO and MBZ both 2D and 3D models show one H-bond with Q134, 167Y and E198. For FBZ we again see H-bond formation with Q134, 167Y and E198, but an additional bond with V236 is also seen in the 2D model. In both models for OXBZ two H-bonds are made with E198 only.

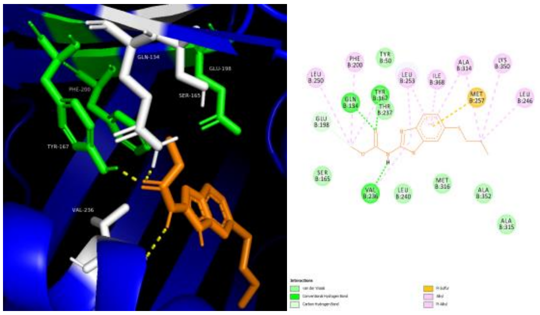

ALA_167Y- ABZ -4.4 kcal/mol

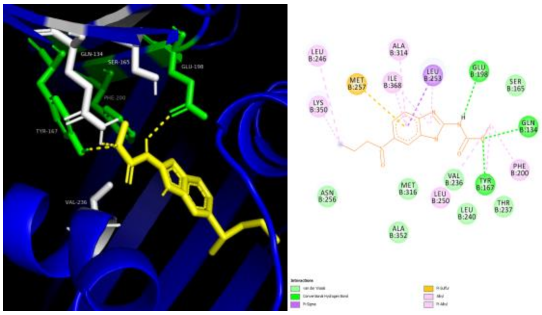

ALA_167Y- ABZSO -4.6

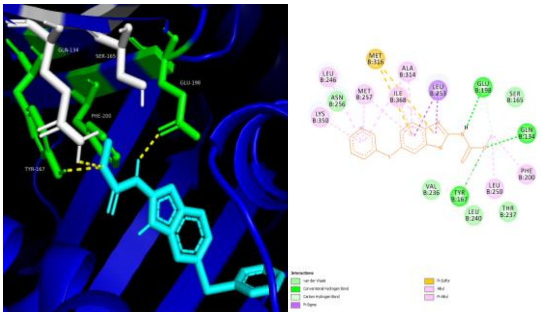

ALA_167Y- FBZ -5 kcal/mol

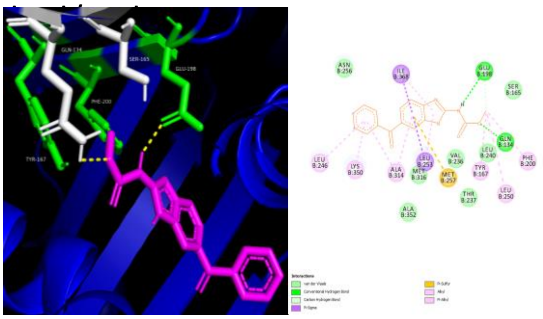

ALA_167Y- MBZ -5.9

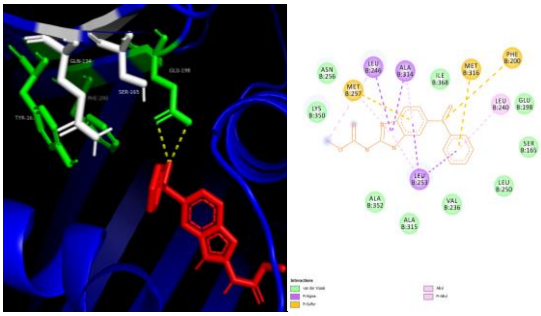

ALA_167Y- OXBZ -3.3 kcal/mol

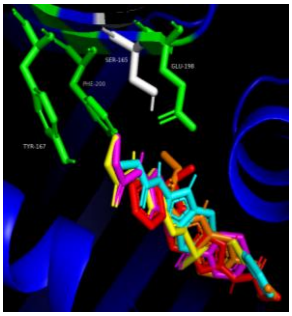

ALA_167Y- all drugs

**Supplementary Figure S5: Autodock vina docking results for *A. lumbricoides* 167Y mutated isotype A and several benzimidazole drugs.** The drugs used are albendazole (ABZ), albendazole sulfoxide (ABZSO), fenbendazole (FBZ), mebendazole (MBZ) and oxfendazole (OXBZ). 3D and 2D models are shown for each docking result. Binding affinities are shown underneath each model. In these mutated 167Y models we see the larger tyrosine amino acid forming bonds with the drugs whereas the natural phenylalanine amino acid does not interact in any other amino acids. In ABZ H-bonds are formed with Q134, 167Y and V236 in both models. For both ABZSO and FBZ H-bonds are made with Q134, 167Y and E198 in 2D and 3D models. In MBZ both models show H-bond formation with Q134 and E198. Finally, in OXBZ two H-bonds are made with E198 in the 3D model but none are predicted in the 2D model.

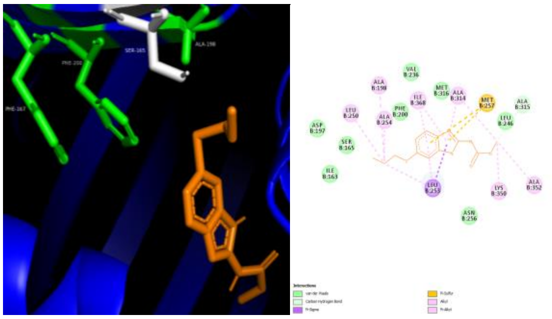

ASA_198A-ABZ -5.7 kcal/mol

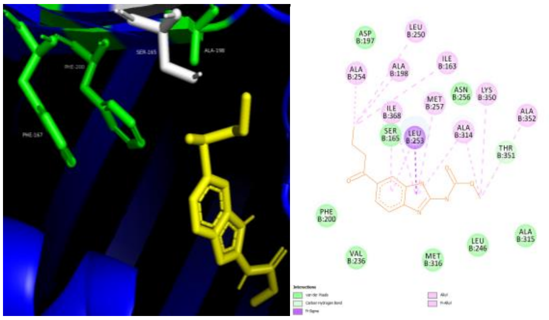

ASA_198A-ABZSO -6.2kcal/mol

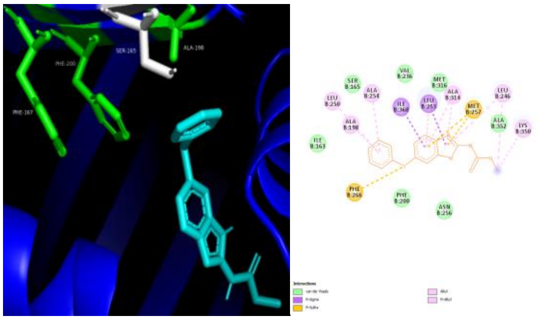

ASA_198A-FBZ -6.2 kcal/mol

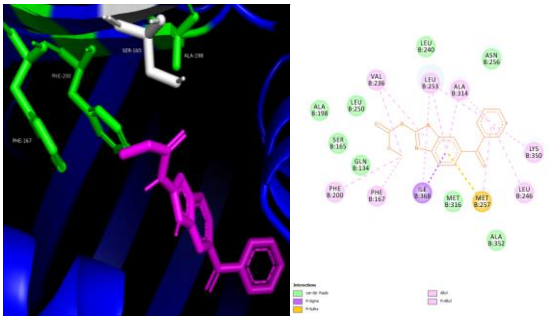

ASA_198A-MBZ -7.6 kcal/mol

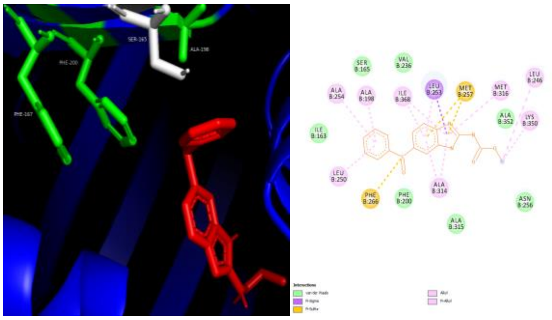

ASA_198A-OXBZ -6.1 kcal/mol

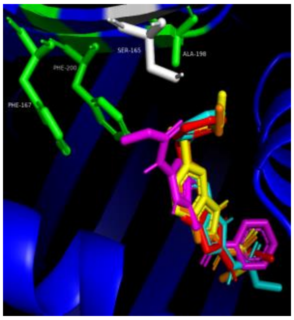

ASA_198A-all drugs

**Supplementary Figure S6: Autodock vina docking results for *A. suum* 198A mutated isotype A and several benzimidazole drugs**. The drugs used are albendazole (ABZ), albendazole sulfoxide (ABZSO), fenbendazole (FBZ), mebendazole (MBZ) and oxfendazole (OXBZ). 3D and 2D models are shown for each docking result. Binding affinities are shown underneath each model. When the mutated 198A amino acid is present, it affects the binding and no H-bond formation is seen with any drug.

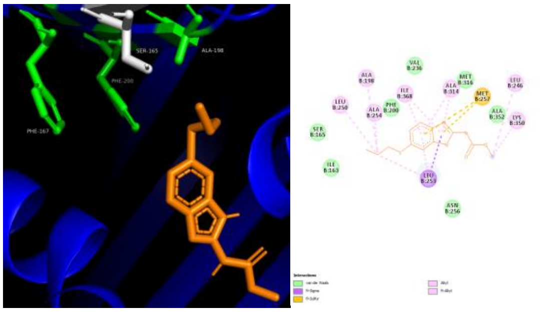

ALA_198A- ABZ -5.7 kcal/mol

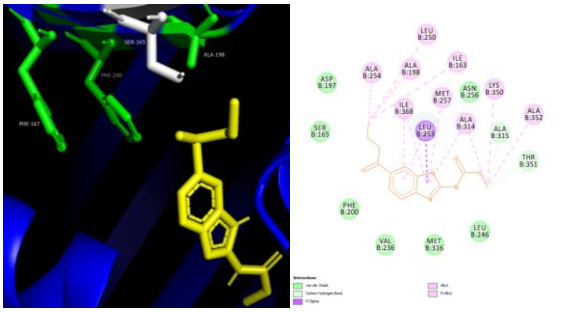

ALA_198A- ABZSO -6.2 kcal/mol

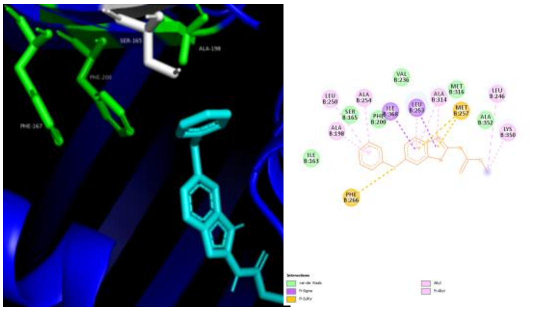

ALA_198A- FBZ -6.2 kcal/mol

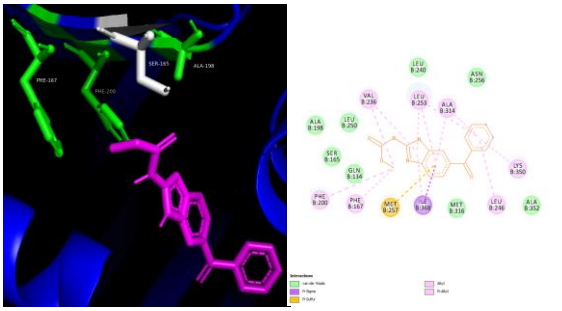

ALA_198A- MBZ -7.6 kcal/mol

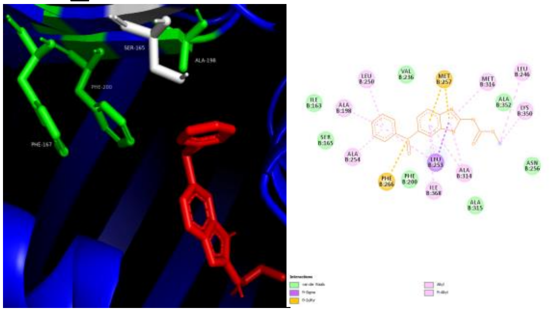

ALA_198A- OXBZ -6.1 kcal/mol

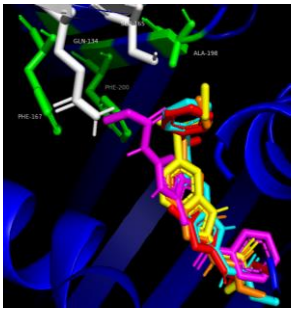

ALA_198A- all drugs

**Supplementary Figure S7: Autodock vina docking results for *A. lumbricoides* 198A mutated isotype A and several benzimidazole drugs**. The drugs used are albendazole (ABZ), albendazole sulfoxide (ABZSO), fenbendazole (FBZ), mebendazole (MBZ) and oxfendazole (OXBZ). 3D and 2D models are shown for each docking result. Binding affinities are shown underneath each model. When the mutated 198A amino acid is present, it affects the binding and no H-bond formation is seen with any drug.

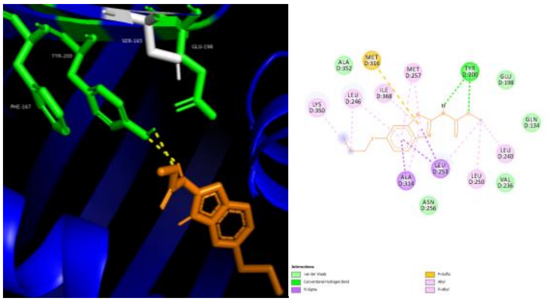

ASA_200Y-ABZ -5.5 kcal/mol

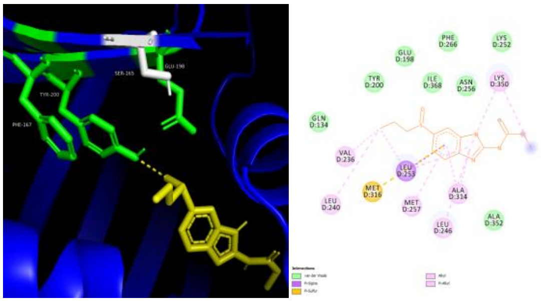

ASA_200Y-ABZSO -5.2

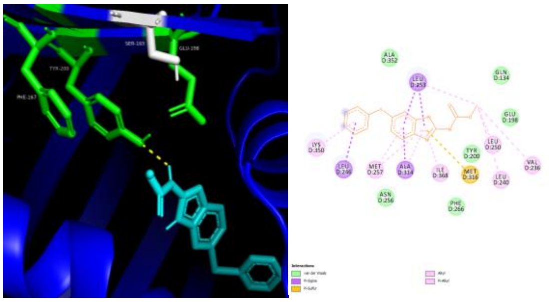

ASA_200Y-FBZ -6 kcal/mol

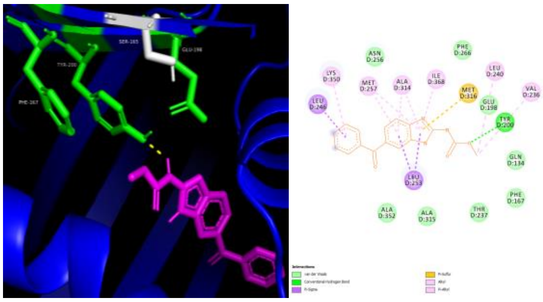

ASA_200Y-MBZ -6.4 kcal/mol

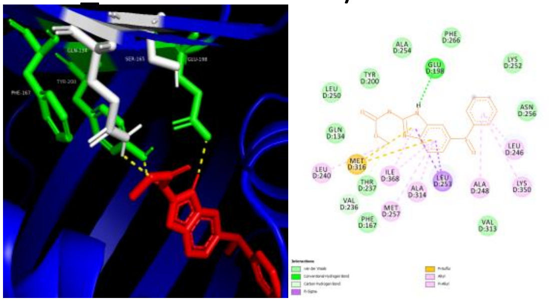

ASA_200Y-OXBZ -4.4 kcal/mol

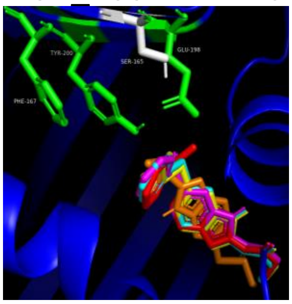

ASA_200Y-all drugs

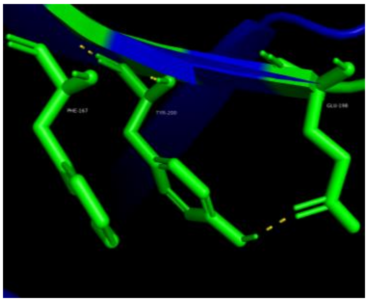

ASA_200Y- Self binding

**Supplementary Figure S8: Autodock vina docking results for *A. suum* 200Y mutated isotype A and several benzimidazole drugs**. The drugs used are albendazole (ABZ), albendazole sulfoxide (ABZSO), fenbendazole (FBZ), mebendazole (MBZ) and oxfendazole (OXBZ). 3D and 2D models are shown for each docking result. Binding affinities are shown underneath each model. When amino acid 200 is mutated to tyrosine the close proximity between the 200Y and E198 amino acids allows H-bonds to form between them. In ABZ both models show two H-bonds with 200Y. For ABZSO and FBZ one H-bond is seen with 200Y in the 3D models but no bonds are seen in the 2D models. In MBZ a H-bond is formed with 200Y in both models. The 3D models for OXBZ show H-bonds with Q134, E198 and 200Y but only the E198 bond is seen in the 2D model.

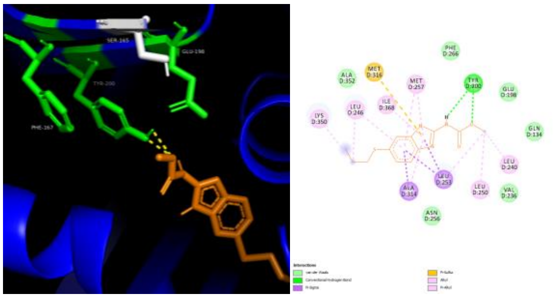

ALA_200Y- ABZ -5.4 kcal/mol

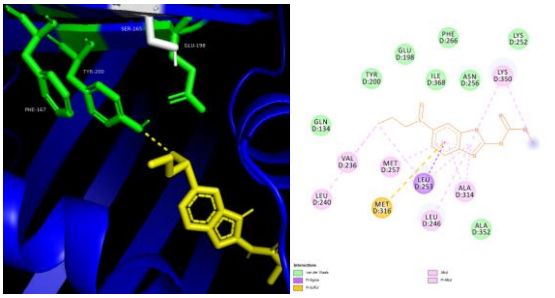

ALA_200Y- ABZSO -5.3 [illegible]

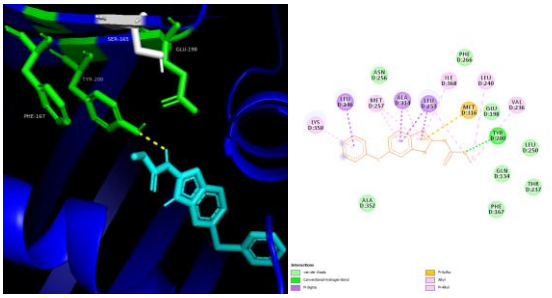

ALA_200Y- FBZ -6 kcal/mol

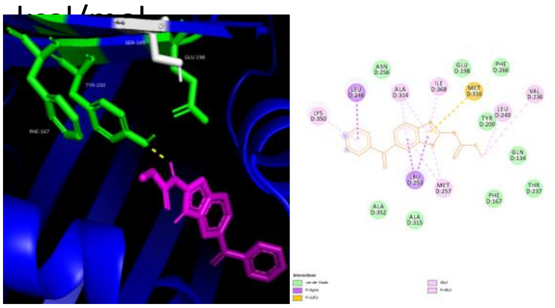

ALA_200Y- MBZ -6.6 [illegible]

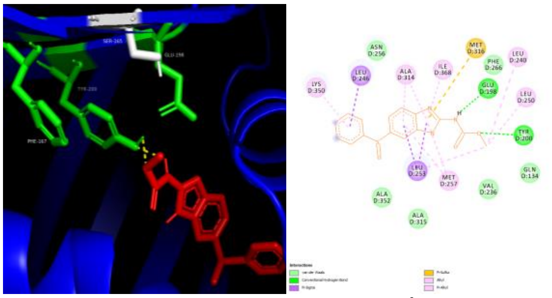

ALA_200Y- OXBZ -4.7 kcal/mol

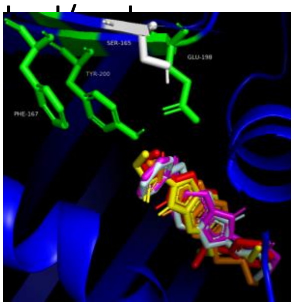

ALA_200Y- all drugs

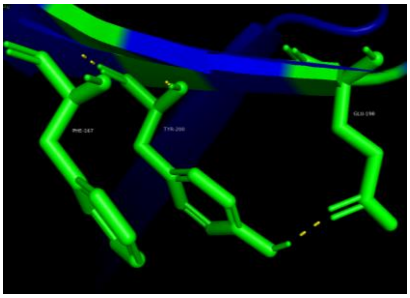

ALA_200Y- self binding

**Supplementary Figure S9: Autodock vina docking results for *A. lumbricoides* 200Y mutated isotype A and several benzimidazole drugs**. The drugs used are albendazole (ABZ), albendazole sulfoxide (ABZSO), fenbendazole (FBZ), mebendazole (MBZ) and oxfendazole (OXBZ). 3D and 2D models are shown for each docking result. Binding affinities are shown underneath each model. When amino acid 200 is mutated to tyrosine the close proximity between the 200Y and E198 amino acids allows H-bonds to form between them. In ABZ both models show two H-bonds with 200Y. For ABZSO and MBZ one H-bond is seen with 200Y in the 3D models but no bonds are seen in the 2D models. In FBZ a H-bond is formed with 200Y in both models. The OXBZ 3D model shows two H-bonds with 200Y but the 2D model shows only one with 200Y and E198.

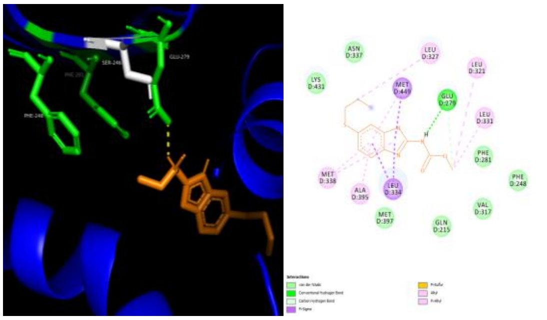

ASB-ABZ -4.3 kcal/mol

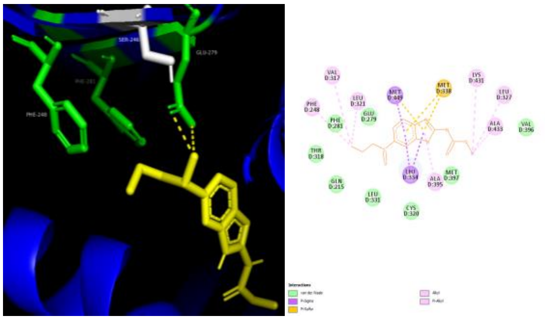

ASB-ABZSO -4.6 kcal/mol

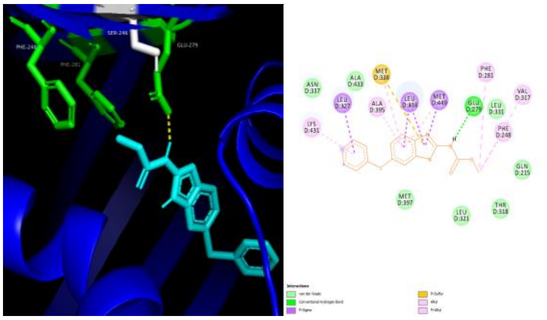

ASB-FBZ -5.1 kcal/mol

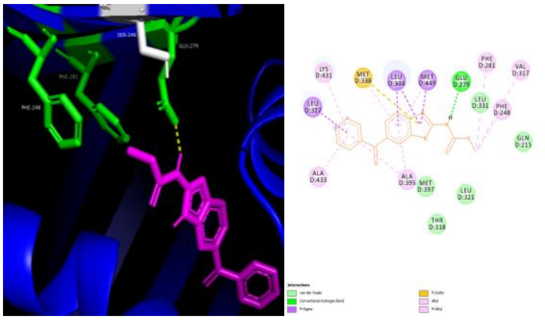

ASB-MBZ -6.5 kcal/mol

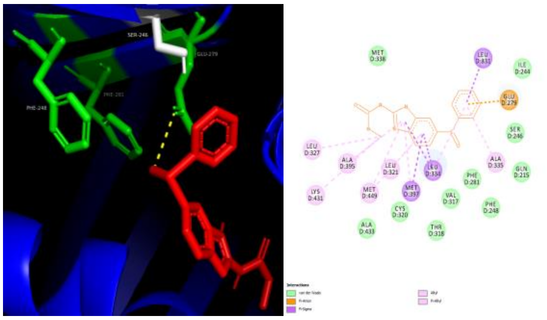

ASB-OXBZ -3.3 kcal/mol

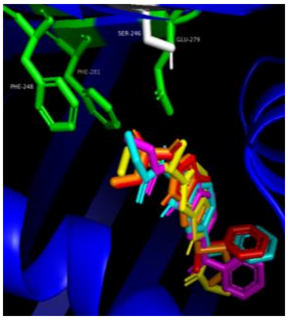

ASB-all drugs

**Supplementary Figure S10: Autodock vina docking results for *A. suum* isotype B and several benzimidazole drugs**. The drugs used are albendazole (ABZ), albendazole sulfoxide (ABZSO), fenbendazole (FBZ), mebendazole (MBZ) and oxfendazole (OXBZ). 3D and 2D models are shown for each docking result. Binding affinities are shown underneath each model. A single H-bond with E198 (labelled as E279 due to isotype B being a longer protein) is seen in all models for each drug, the exception is that for the ABZSO and OXBZ 2D models no H-bonds are found.

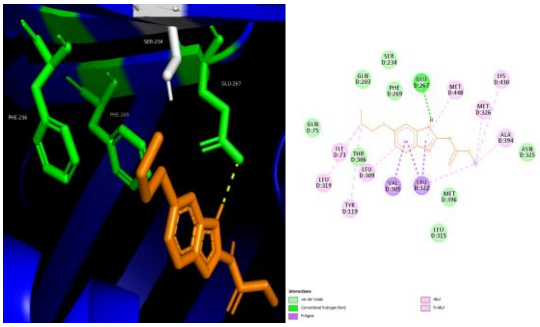

ALB- ABZ -5.9 kcal/mol

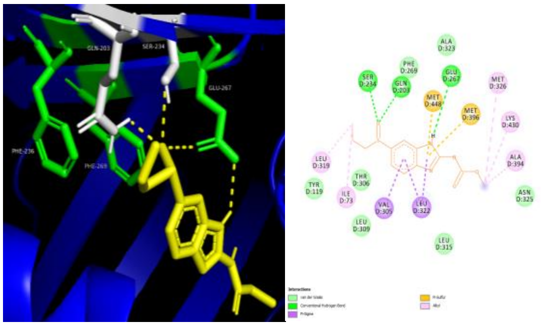

ALB- ABZSO -6.1 kcal/mol

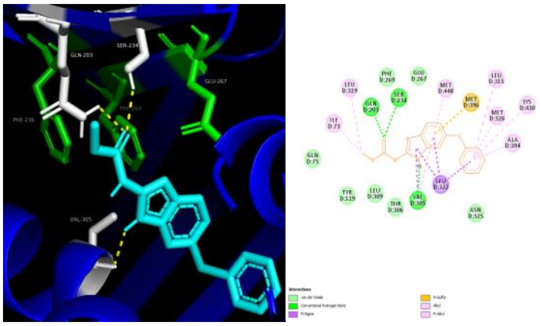

ALB- FBZ -7.3 kcal/mol

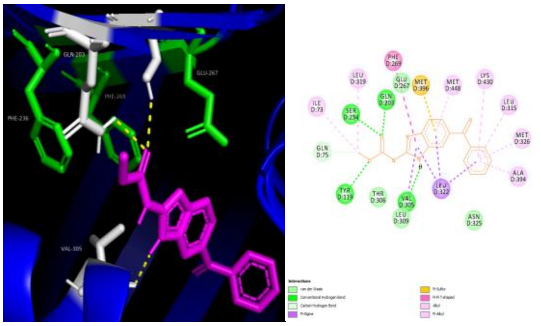

ALB- MBZ -8.5 kcal/mol

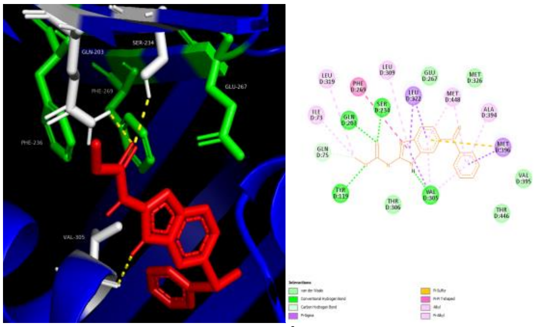

ALB- OXBZ -6 kcal/mol

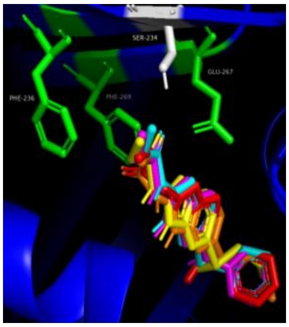

ALB- all drugs

**Supplementary Figure S11: Autodock vina docking results for *A. lumbricoides* isotype B and several benzimidazole drugs**. The drugs used are albendazole (ABZ), albendazole sulfoxide (ABZSO), fenbendazole (FBZ), mebendazole (MBZ) and oxfendazole (OXBZ). 3D and 2D models are shown for each docking result. Binding affinities are shown underneath each model. For ABZ a single H-bond with E198 (labelled as E267 due to isotype B being a longer protein) is seen in both 2D and 3D models. For ABZSO we find two H-bonds with E198, one with Q134 (Q203) and one with S165 (S234) in the 3D model, with only one bond seen to each in the 2D model. For FBZ, MBZ and OXBZ the models show H-bonds to Q134, S165 and V236 (V305) in all models, with the addition of Y50 (Y119) in the 2D models for MBZ and OXBZ.

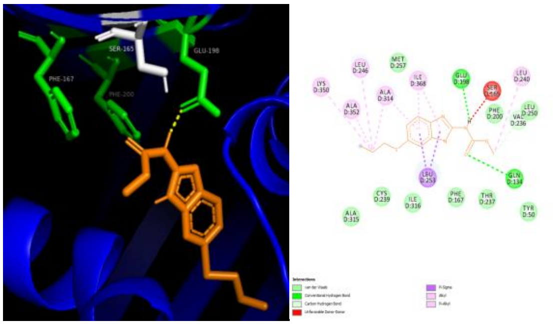

ASC-ABZ -6.6 kcal/mol

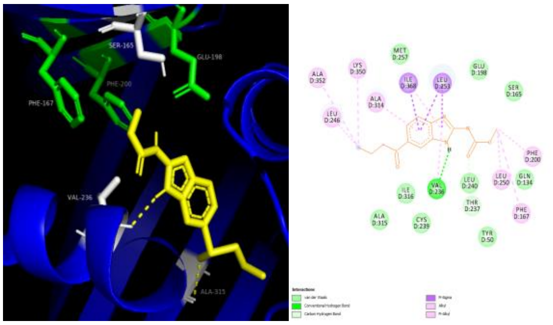

ASC-ABZSO -6.6 kcal/mol

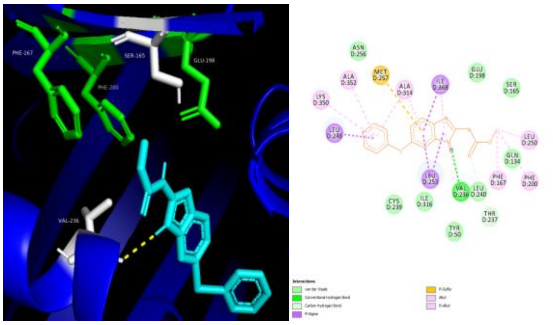

ASC-FBZ -7.8 kcal/mol

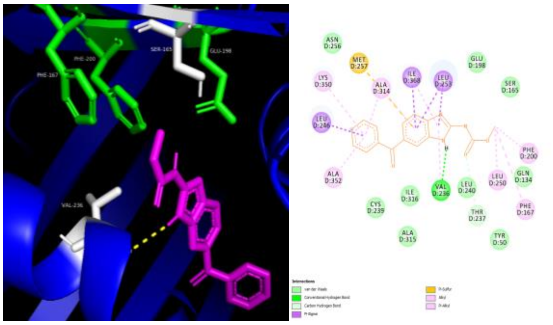

ASC-MBZ -8.4 kcal/mol

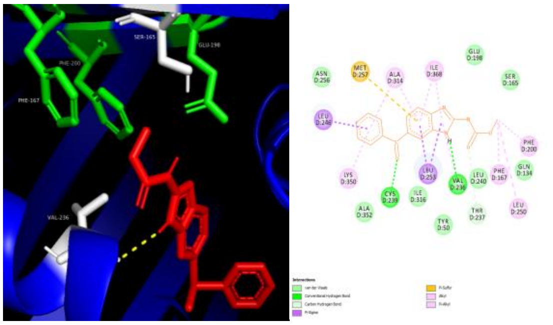

ASC-OXBZ -7.2 kcal/mol

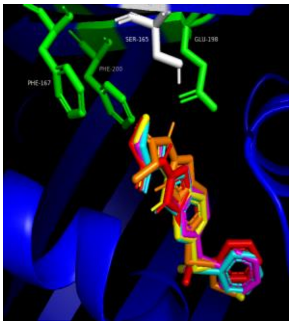

ASC-all drugs

**Supplementary Figure S12: Autodock vina docking results for *A. suum* isotype C and several benzimidazole drugs**. The drugs used are albendazole (ABZ), albendazole sulfoxide (ABZSO), fenbendazole (FBZ), mebendazole (MBZ) and oxfendazole (OXBZ). 3D and 2D models are shown for each docking result. Binding affinities are shown underneath each model. In ABZ 3D docking models we find a H-bond formed with E198, yet in the 2D models we see that on top of this there is also a bond with Q134 and although unfavourable there is a donor-donor bond with S165. For all other drugs we see a single H-bond formed with V236 in both models, with an extra bond to A315 in ABZSO 3D model and C239 in the OXBZ 2D model.

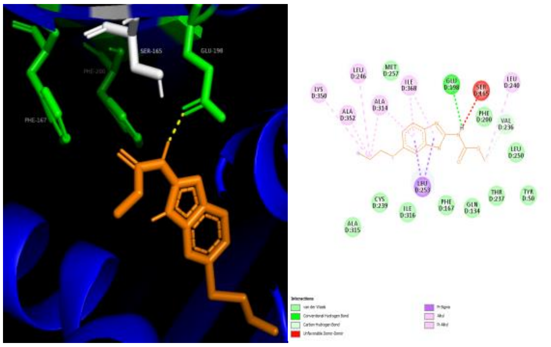

ALC- ABZ -6.6 kcal/mol

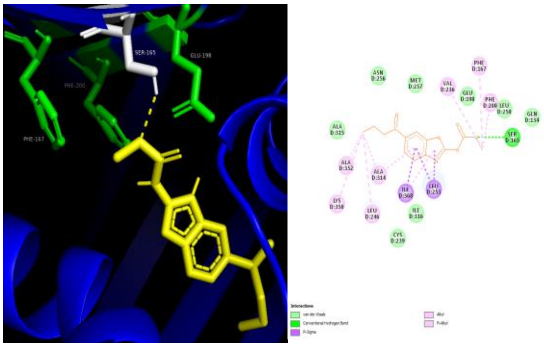

ALC- ABZSO -6.1 kcal/mol

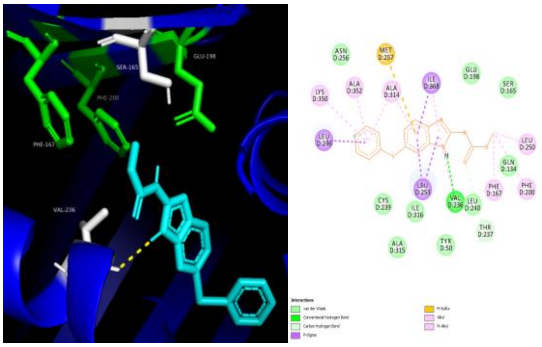

ALC- FBZ -7.8 kcal/mol

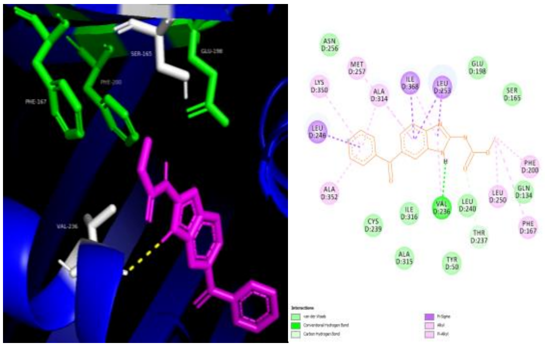

ALC- MBZ -8.5 kcal/mol

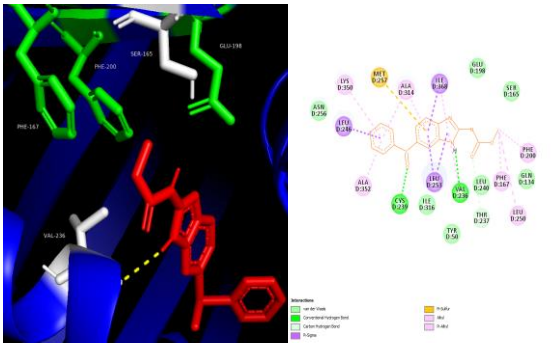

ALC- OXBZ -7.2 kcal/mol

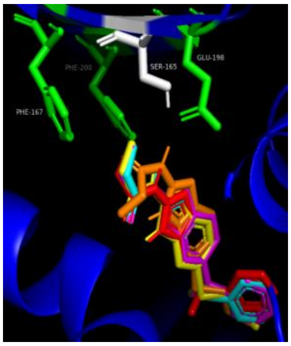

ALC- all drugs

**Supplementary Figure S13: Autodock vina docking results for *A. lumbricoides* isotype C and several benzimidazole drugs.** The drugs used are albendazole (ABZ), albendazole sulfoxide (ABZSO), fenbendazole (FBZ), mebendazole (MBZ) and oxfendazole (OXBZ). 3D and 2D models are shown for each docking result. Binding affinities are shown underneath each model. In ABZ 3D docking models we find a H-bond formed with E198, yet in the 2D models we see that on top of this there is also a bond with Q134 and although unfavourable there is a donor-donor bond with S165. In the ABZSO 3D and 2D docking models one H-bond is formed with S165 only. For all other drugs we see a single H-bond formed with V236 in both models, with an extra bond to A315 in ABZSO 3D model and C239 in the OXBZ 2D model.

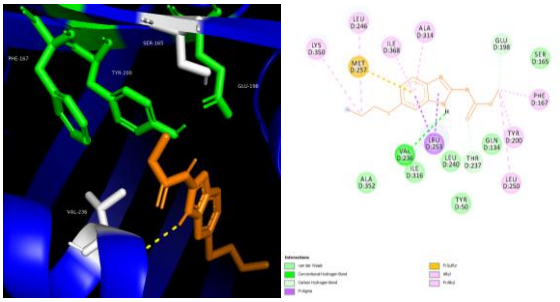

ASD-ABZ -6.2 kcal/mol

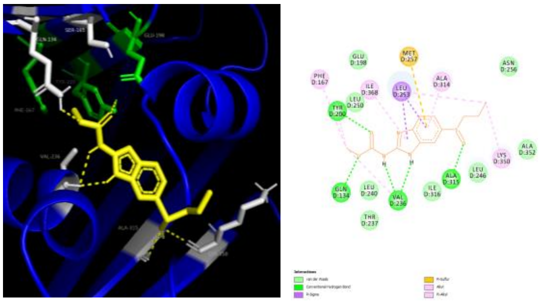

ASD-ABZSO -6.8 kcal/mol

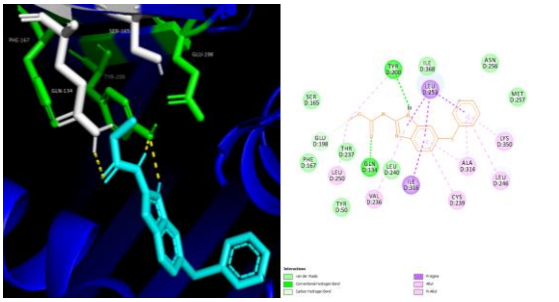

ASD-FBZ -7.1 kcal/mol

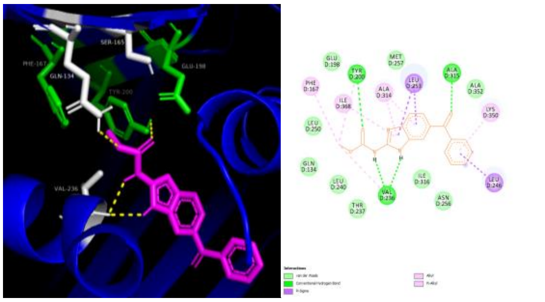

ASD-MBZ -8.6 kcal/mol

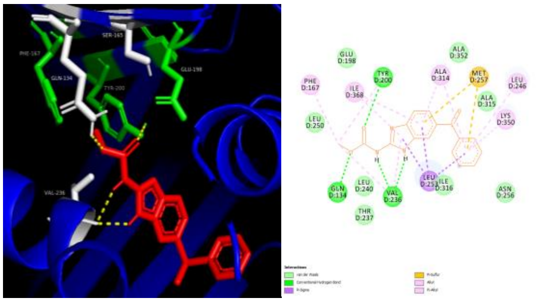

ASD-OXBZ -6.3 kcal/mol

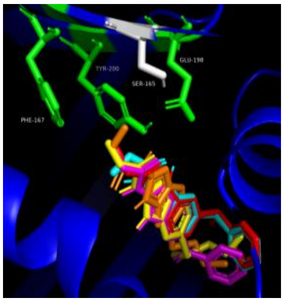

ASD-all drugs

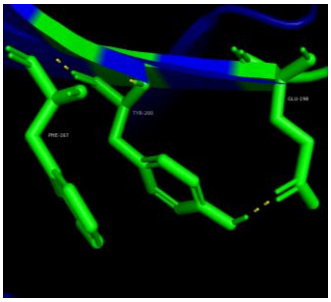

ASD-self binding

**Supplementary Figure S14: Autodock vina docking results for *A. suum* isotype D and several benzimidazole drugs.** The drugs used are albendazole (ABZ), albendazole sulfoxide (ABZSO), fenbendazole (FBZ), mebendazole (MBZ) and oxfendazole (OXBZ). 3D and 2D models are shown for each docking result. Binding affinities are shown underneath each model. A single H-bond is made with V236 and ABZ in both 3D and 2D models. For ABZSO we see two H-bonds with amino acids V236 and A315 and one H-bond with Q134, Y200 and K350 in the 3D model. In the 2D model we again find two H-bonds with V236, but only a single bond with A315, along with Q134 and Y200. In the FBZ models we see one H-bond made with Q134 in both 3D and 2D model, H-bonds are also formed with Y200 in both models with two seen in the 3D and one in the 2D models. For MBZ we see the formation of two H-bonds to V236 and one with Y200 in both models, in addition to this the 3D model shows an extra bond with Q134 and the 2D model shows a bond with A315. Finally, in the OXBZ models we again find the formation of two H-bonds with V236, along with single bonds to Q134 and Y200 in both models.

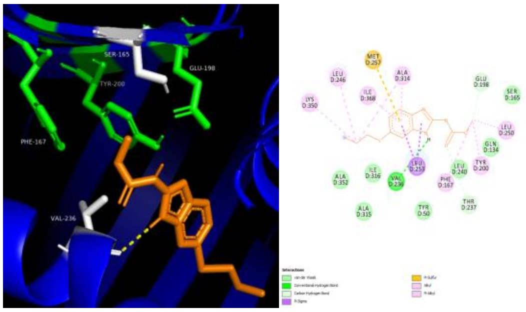

ALD- ABZ -6.2 kcal/mol

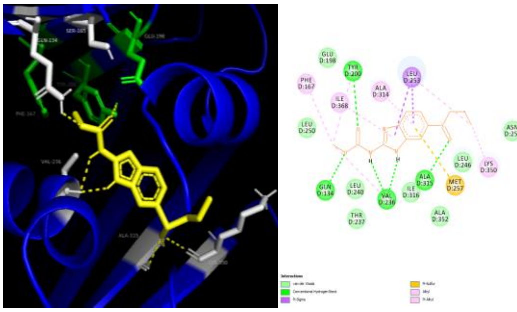

ALD- ABZSO -6.8 kcal/mol

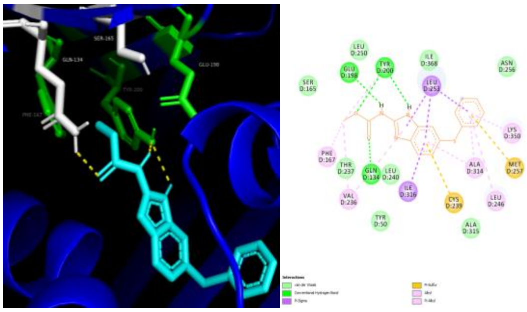

ALD- FBZ -7 kcal/mol

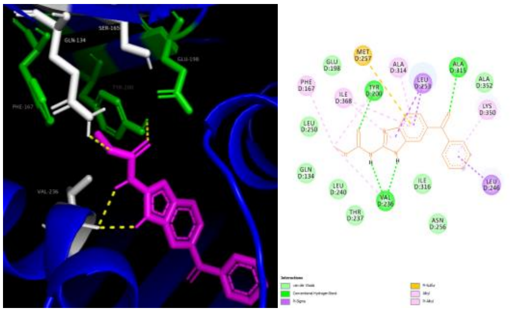

ALD- MBZ -8.5 kcal/mol

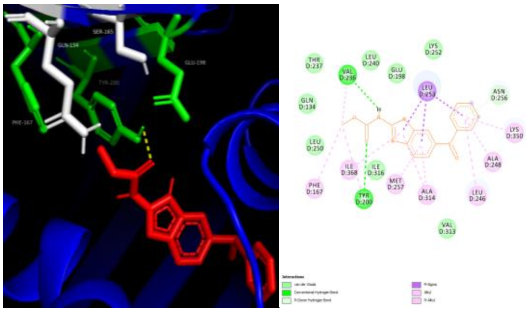

ALD- OXBZ -7.1 kcal/mol

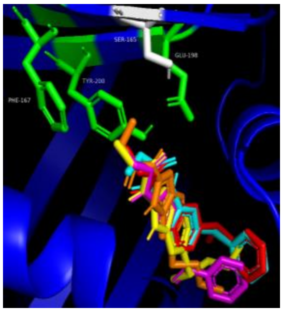

ALD- all drugs

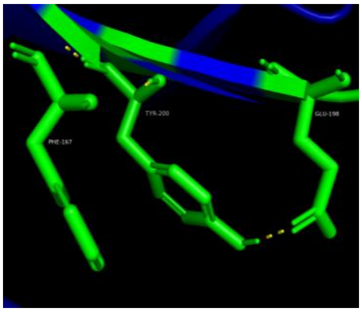

ALD- self binding

**Supplementary Figure S15: Autodock vina docking results for *A. lumbricoides* isotype D and several benzimidazole drugs.** The drugs used are albendazole (ABZ), albendazole sulfoxide (ABZSO), fenbendazole (FBZ), mebendazole (MBZ) and oxfendazole (OXBZ). 3D and 2D models are shown for each docking result. Binding affinities are shown underneath each model. For ABZ models one H-bond is formed with Q134 in both models. The 3D model then shows a H-bond with E198, whilst the 2D models show two bonds between the drug and E198. For ABZSO we see two H-bonds with amino acids V236 and A315 and one H-bond with Q134, Y200 and K350 in the 3D model. In the 2D model we again find two H-bonds with V236, but only a single bond with A315, along with Q134 and Y200. With FBZ two H-bonds form with Y200 and one forms with Q134 in both models, there is also an additional bond with E198 seen in the 2D model.  For MBZ we see the formation of two H-bonds to V236 and one with Y200 in both models, in addition to this the 3D model shows an extra bond with Q134 and the 2D model shows a bond with A315. The OXBZ models both show a single H-bond formation with Y200, however the 2D models show an additional bond with V236.

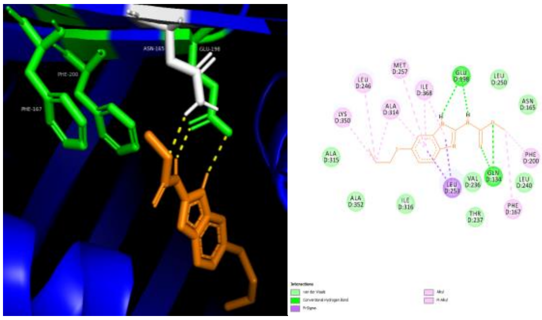

ASE- ABZ -5.6 kcal/mol

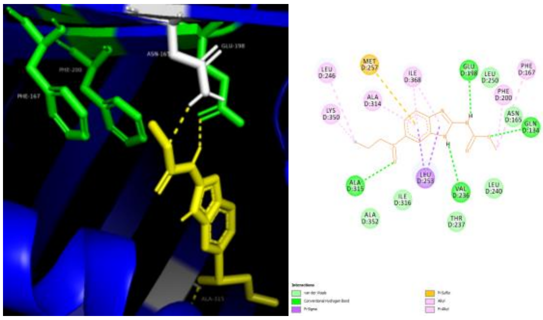

ASE- ABZSO -5.9 kcal/mol

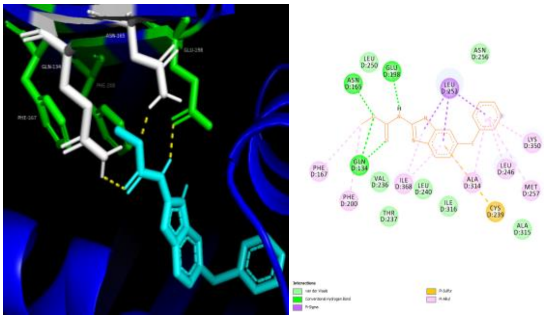

ASE- FBZ -6.9 kcal/mol

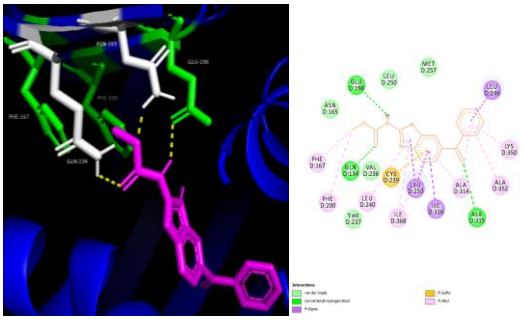

ASE- MBZ -7.4 kcal/mol

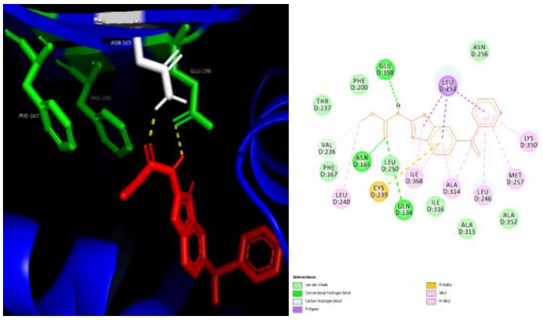

ASE- OXBZ -6.5 kcal/mol

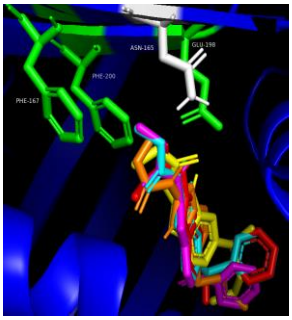

ASE- all drugs

**Supplementary Figure S16: Autodock vina docking results for *A. suum* isotype E and several benzimidazole drugs.** The drugs used are albendazole (ABZ), albendazole sulfoxide (ABZSO), fenbendazole (FBZ), mebendazole (MBZ) and oxfendazole (OXBZ). 3D and 2D models are shown for each docking result. Binding affinities are shown underneath each model. In ABZ, docking shows two H-bonds with E198 and one with N165 in the 3D model but two H-bonds with both Q134 and E198 in the 2D model. With ABZSO the 3D models show one H-bond with N165 and E198, however the 2D model predicts H-bonds with Q134, E198, V236 and A315. Both models for FBZ find single H-bonds with Q134, N165 and E198. In the MBZ 3D model one H-bond is predicted to form with Q134, N165 and E198 whereas in the 2D model the H-bonds are seen with Q134, E198 and A315. Finally, the OXBZ 3D model predicts H-bonds with N165 and E198, whilst the 2D model shows an additional bond with Q134.

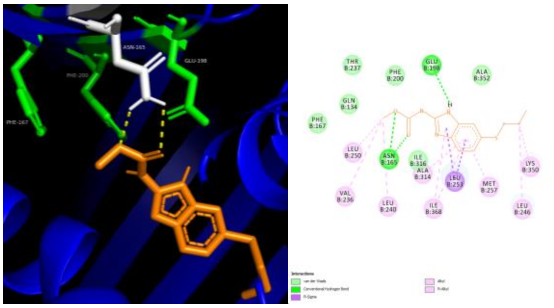

ALE- ABZ -6.1 kcal/mol

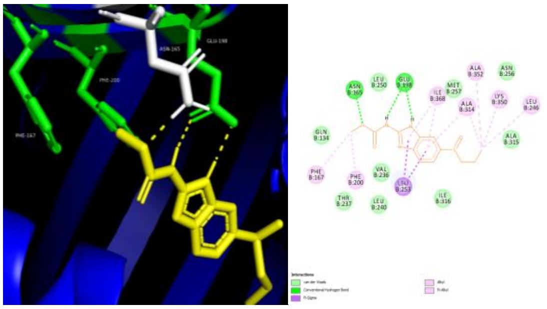

ALE- ABZSO -6.8 kcal/mol

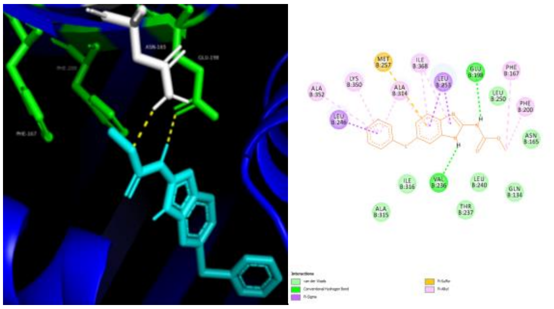

ALE- FBZ -7.5 kcal/mol

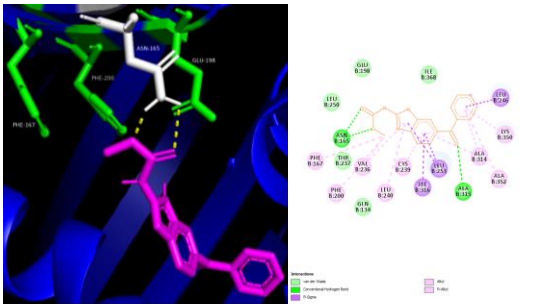

ALE- MBZ -8.3 kcal/mol

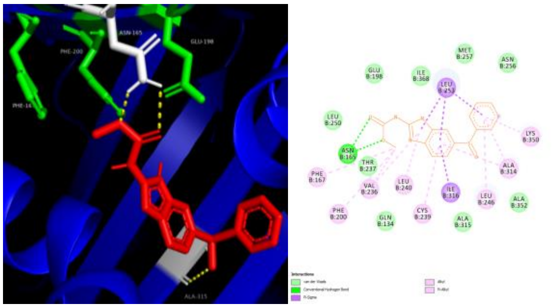

ALE- OXBZ -7.3 kcal/mol

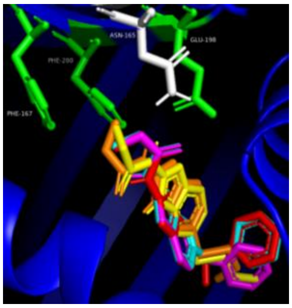

ALE- all drugs

**Supplementary Figure S17: Autodock vina docking results for *A. lumbricoides* isotype E and several benzimidazole drugs.** The drugs used are albendazole (ABZ), albendazole sulfoxide (ABZSO), fenbendazole (FBZ), mebendazole (MBZ) and oxfendazole (OXBZ). 3D and 2D models are shown for each docking result. Binding affinities are shown underneath each model. In ABZ docking two H-bonds are seen with N165 in the 3D model with an additional bond with E198 shown in the 2D model.  for ABZSO both models predict the formation of one H-bond to N165 and two H-bonds to E198. For FBZ H-bond formation with N165 and E198 is seen in the 3D model, whereas H-bonds are made with E198 and V236 in the 2D model. In the 3D model for MBZ two H-bonds are made with N165 and in the 2D model an additional bond with A315 is seen. Finally, for OXBZ the 3D model shows two H-bonds with N165 and one with A315, however the 2D model only finds the two bonds with N165.

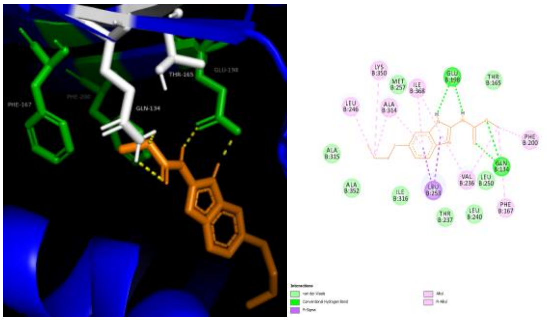

ASF- ABZ -7.2 kcal/mol

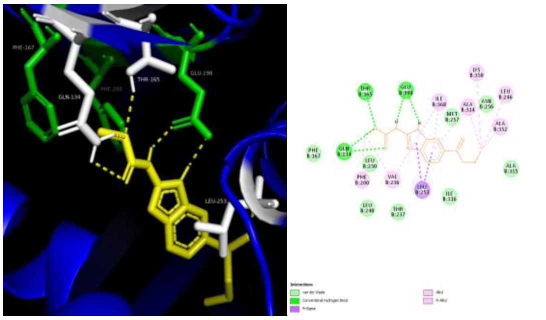

ASF- ABZSO -7.7 kcal/mol

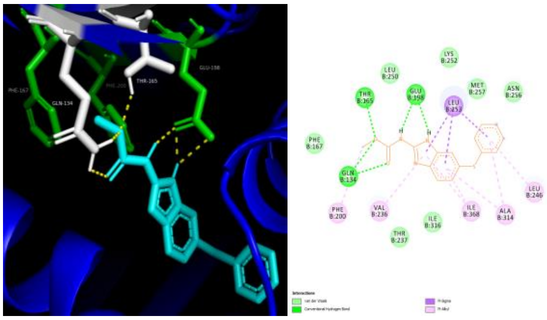

ASF- FBZ -7.5 kcal/mol

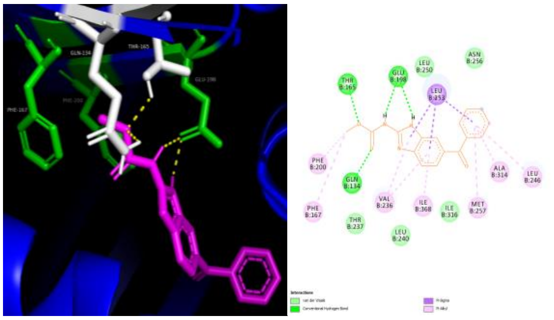

ASF- MBZ -8.4 kcal/mol

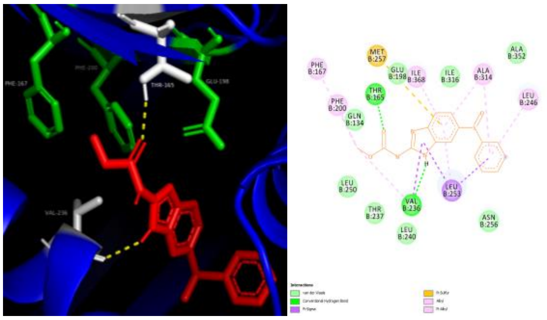

ASF- OXBZ -7.1 kcal/mol

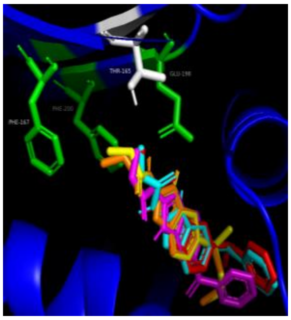

ASF- all drugs

**Supplementary Figure S18: Autodock vina docking results for *A. suum* isotype F and several benzimidazole drugs.** The drugs used are albendazole (ABZ), albendazole sulfoxide (ABZSO), fenbendazole (FBZ), mebendazole (MBZ) and oxfendazole (OXBZ). 3D and 2D models are shown for each docking result. Binding affinities are shown underneath each model. In ABZ both docking models predict two H-bonds with both Q134 and E198. For ABZSO two H-bonds are made with Q134 and E198 and a single bond is formed with T165 in both 3D and 2D models. The FBZ 3D model predicts three H-bonds with E198, two H-bonds with Q 134 and one bond with T165, the 2D model is similar but only shows two H-bonds with E198. In both models for MBZ two H-bonds are seen with E198 and one is seen with Q134 and T165. The final drug, OXBZ has H-bonds with T165 and V236 in both models.

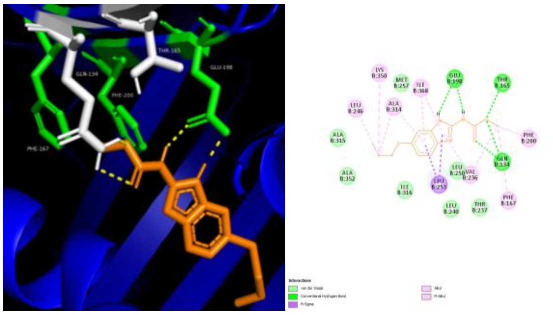

ALF- ABZ -7.2 kcal/mol

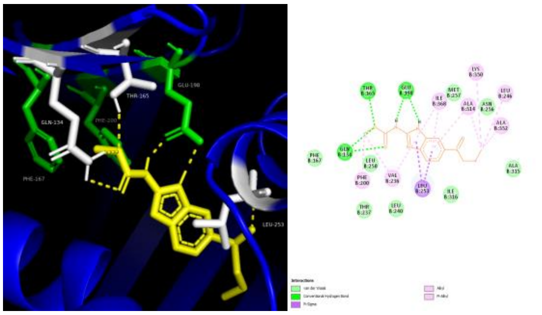

ALF- ABZSO -7.8 kcal/mol

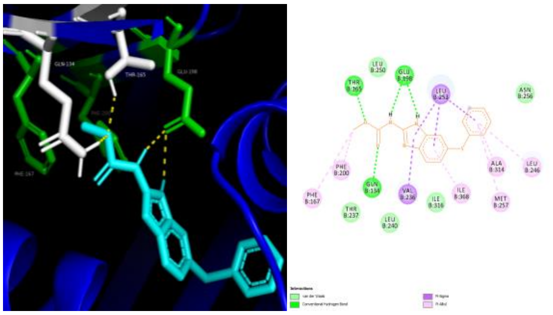

ALF- FBZ -7.4 kcal/mol

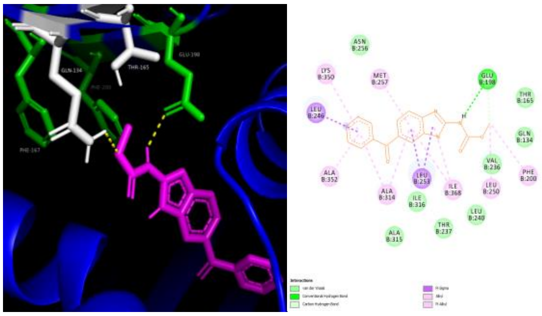

ALF- MBZ -8.7 kcal/mol

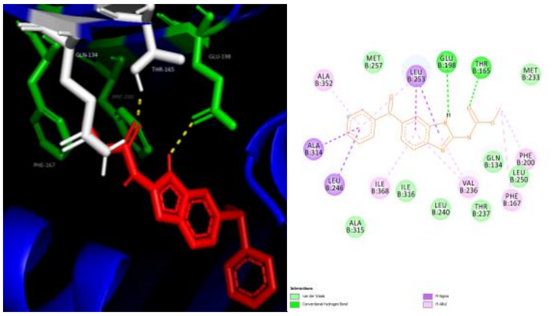

ALF- OXBZ -6 kcal/mol

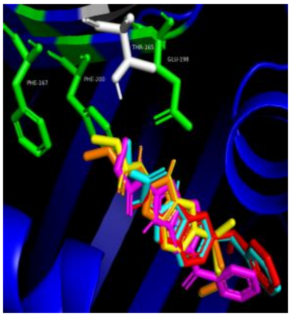

ALF- all drugs

**Supplementary Figure S19: Autodock vina docking results for *A. lumbricoides* isotype F and several benzimidazole drugs.** The drugs used are albendazole (ABZ), albendazole sulfoxide (ABZSO), fenbendazole (FBZ), mebendazole (MBZ) and oxfendazole (OXBZ). 3D and 2D models are shown for each docking result. Binding affinities are shown underneath each model. In ABZ both models show two H-bonds with both Q134 and E198 but in the 2D model an extra H-bond with T165 is also shown. For ABZSO two H-bonds are shown with Q134 and E198 and one H-bond is formed with T165 and L253 in the 3D model, however in the 2D model the bond with L253 is not found. In FBZ two H-bonds are formed with E198 and one bond is formed with Q143 and T165 in both models. The 3D model for MBZ shows single H-bonds with Q134 and E198 however the 2D model shows only the bond with E198. Finally, both models for OXBZ predict H-bond formation with T165 and E198.

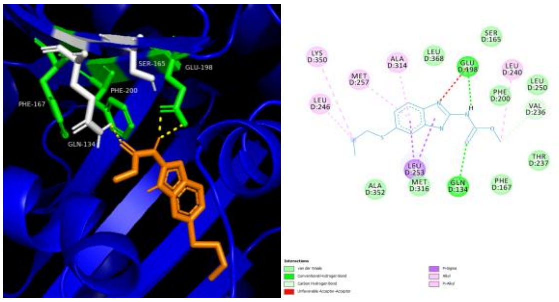

ASG- ABZ-4.1 kcal/mol

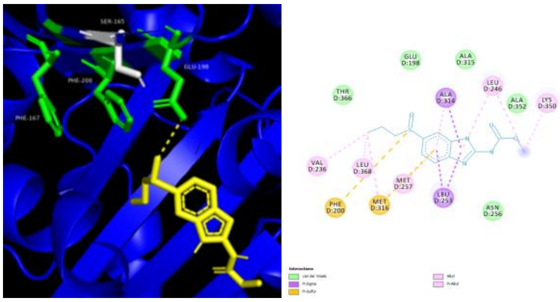

ASG- ABZSO -4.1 kcal/mol

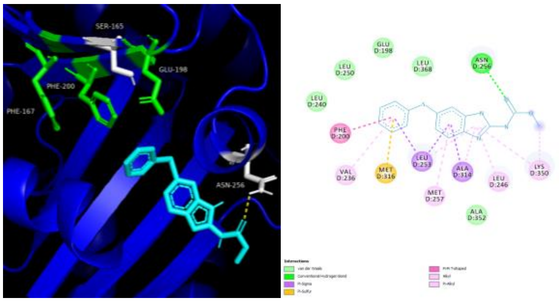

ASG- FBZ -6.2 kcal/mol

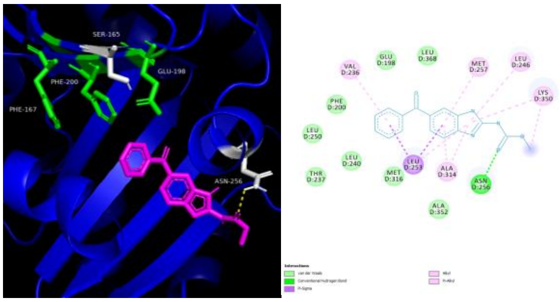

ASG- MBZ -6.9 kcal/mol

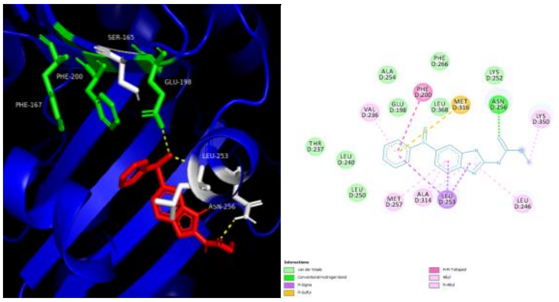

ASG- OXBZ -5.2 kcal/mol

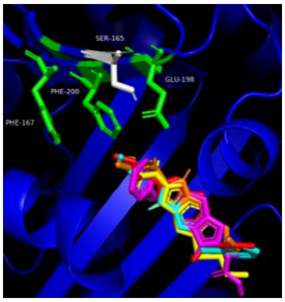

ASG- all drugs

**Supplementary Figure S20: Autodock vina docking results for A. suum isotype G and several benzimidazole drugs.** The drugs used are albendazole (ABZ), albendazole sulfoxide (ABZSO), fenbendazole (FBZ), mebendazole (MBZ) and oxfendazole (OXBZ). 3D and 2D models are shown for each docking result. Binding affinities are shown underneath each model. The interactions between ASG and ABZ form 2 H-bonds with E198 and 1 with Q134 in both 2D and 3D models, however one of the H-bonds with E198 is predicted to be an unfavourable acceptor to acceptor bond in the 2D model. For ABZSO a single H-bond is formed with E198 in the 3D model but no bonds are seen in the 2D model. In the cases of FBZ and MBZ a single H-bond to N265 is seen in both 3D and 2D models. Finally for OXBZ 1 H-bond is formed with E198, L253 and N256 in the 3D model but only a single H-bond is predicted with N256 in the 2D model.

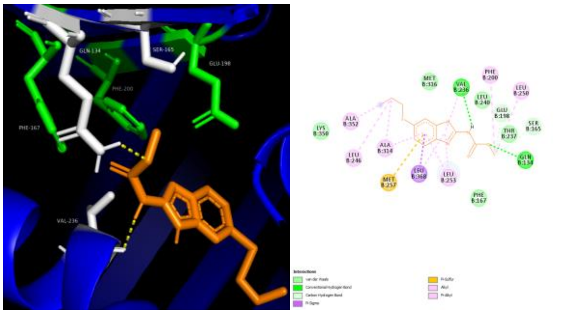

ALG- ABZ -6.2 kcal/mol

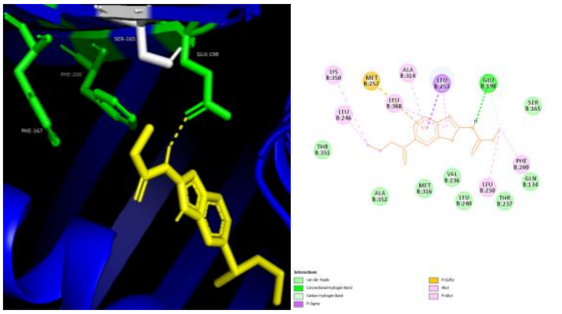

ALG- ABZSO -6.1 kcal/mol

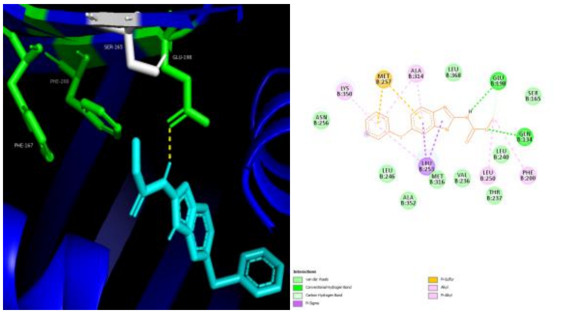

ALG- FBZ -7 kcal/mol

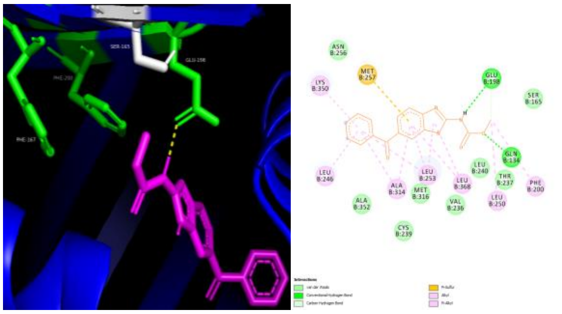

ALG- MBZ -7.8 kcal/mol

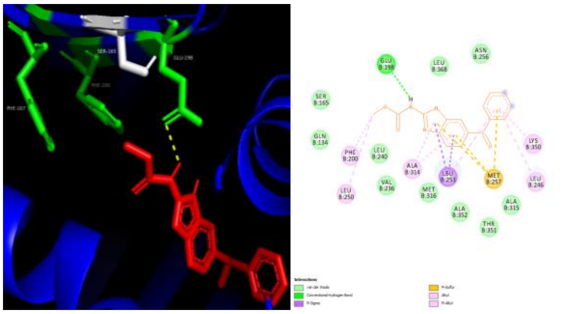

ALG- OXBZ -5.9 kcal/mol

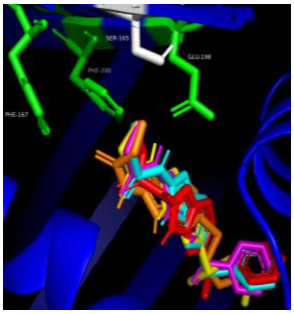

ALG- all drugs

**Supplementary Figure S21: Autodock vina docking results for *A. lumbricoides* isotype L and several benzimidazole drugs.** The drugs used are albendazole (ABZ), albendazole sulfoxide (ABZSO), fenbendazole (FBZ), mebendazole (MBZ) and oxfendazole (OXBZ). 3D and 2D models are shown for each docking result. Binding affinities are shown underneath each model. In ABZ both models find H-bonds with Q134 and V236. Both models for ABZSO and OXBZ show H-bonding with only E198. For FBZ and MBZ the 3D models show H-bonds with E198, whereas the 2D models find H-bond formation with Q134 and E198.
